# Bacterial extracellular vesicles as a tunable platform for vaginal drug delivery

**DOI:** 10.1101/2025.06.09.658669

**Authors:** Darby Steinman, Varunaa Sri Hemanth Kumar, Ryan A McIlvaine, Pranshu Tyagi, Hahnbit Kang, Raifah Alam, Anguo Liu, Christopher M Jewell, Irina Burd, Sara Molinari, Hannah C Zierden

## Abstract

There is a critical gap in the development of new therapeutic platforms designed to treat gynecologic and obstetric diseases. Compared to systemic drug delivery, vaginal drug administration of nanoparticle formulations limits off-target side effects while increasing therapeutic concentration in target tissues, showing promise for clinical translation. However, these formulations suffer from limited scalability, high-cost reagents, and long optimization timelines. Recent work highlights the potential of bacterial extracellular vesicles (bEVs) as a low-cost, tunable platform for therapeutic applications. Here, we evaluate bEVs as a therapeutic carrier for vaginal drug delivery. We demonstrate the loading of the model protein moxNeonGreen into *Escherichia coli* Nissle 1917 bEVs. By optimizing growth parameters, we increase protein loading into bEVs. We evaluate the effect of bEVs on the vaginal microenvironment, and observe no negative impact on vaginal epithelial cells, endocervical cells, or vaginal bacteria *in vitro*. Additionally, we observe the retention of bEVs in the murine female reproductive tract for more than six hours. This study provides a framework for using genetically engineered bEVs to rapidly generate customizable therapies for a range of gynecologic and obstetric conditions, addressing longstanding challenges in women’s health therapeutics.

**Graphical Abstract:** 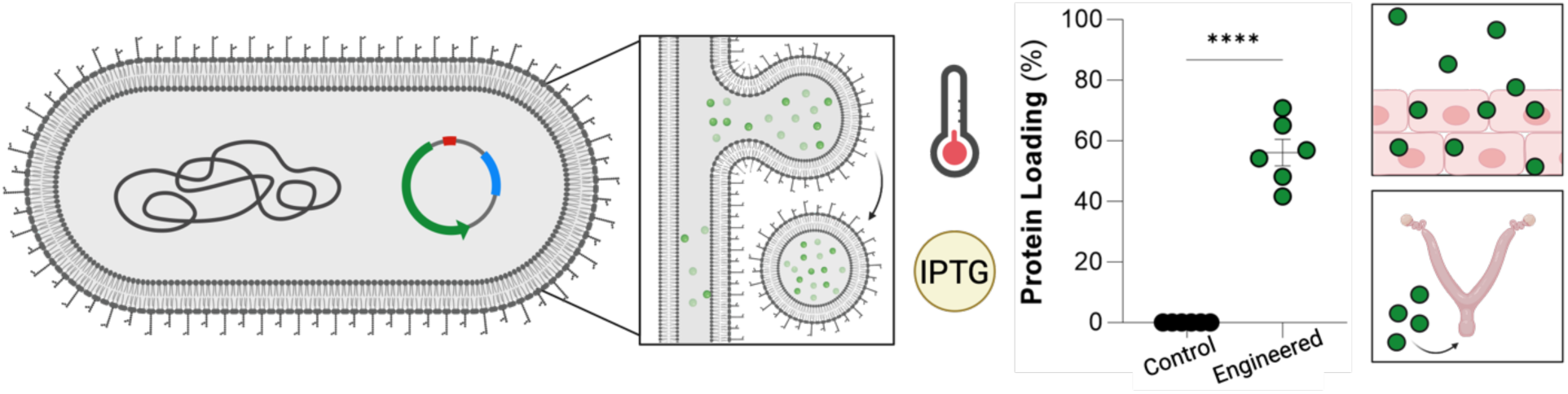

## Introduction

There is a critical need for rapid clinical translation of new therapeutic options aimed at treating gynecologic and obstetric disease. Recent evidence demonstrates that only 3.7% of clinical trials focus on gynecology, with chronic underfunding of research into women’s health indications.^1,2^ In recent years, nanoparticle formulations have had a considerable impact on cancer, immune disorders, and infectious diseases, however, there have been no clinically approved nanoparticle formulations for treating female reproductive health.^3-8^

A growing body of work suggests that vaginally administered nanoparticles are a particularly useful platform for targeting the female reproductive tract. Vaginally administered nanoparticles demonstrate enhanced uptake into target tissues, as well as reduced off-target side-effects, compared to systemic drug delivery. Vaginal drug delivery leads to improved therapeutic outcomes across a range of reproductive diseases, including preterm birth, vaginal atrophy, sexually transmitted infections, and cervical cancer.^9-16^ Despite promising preclinical results, these formulations suffer from complex production parameters with limited scalability, high-cost reagents, and long optimization timelines to expand these nanoparticle platforms to other drug targets. In contrast, recent work highlights the potential of bacterial extracellular vesicles to overcome these manufacturing challenges and serve as a drug delivery platform for a range of human diseases.^10,17-22^

Bacterial extracellular vesicles (bEVs) are membrane-bound particles produced by both Gram-positive and Gram-negative bacteria.^23^ bEVs are typically 100-500 nm in size and reflect the activity of parent cells via their small molecule, protein, and nucleic acid cargoes.^24,25^ Given the relatively low cost of bacterial culture, the adaptability to biomanufacturing infrastructure, and the tunability of therapeutic properties through established bioengineering approaches, bEVs have garnered interest in drug delivery applications, including vaccination, immunotherapy, and antibacterial agents for a range of disease indications.^17,26,27^ For the first time, our work evaluates bEVs as a platform for vaginal drug delivery applications. Using a synthetic biology toolbox, we demonstrate the usefulness of genetic engineering in loading bEVs with target proteins, examine the compatibility of bEVs within the vaginal microenvironment, and demonstrate retention after vaginal administration in a preclinical animal model. These data demonstrate the potential use of bEVs as a pathway to expedite the formulation of new nanomedicines for gynecologic and obstetric indications.

## Materials and Methods

### Bacterial strains

*Escherichia coli* Nissle 1917 (NCBI:txid316435) was genetically modified to produce bEVs throughout these experiments. ^28^ A human-derived strain of *Lactobacillus crispatus* (Brygoo and Aladame, 33820) was sourced from American Type Culture Collection (ATCC; Manassas, VA).

### Human female reproductive tract cell lines

Vaginal epithelial (VK2/E6E7, CRL-2616) and endocervical (End1/E6E7, CRL-2615) cells were sourced from American Type Culture Collection (ATCC; Manassas, VA).

### Materials

Keratinocyte-serum free medium, keratinocyte-serum free media without calcium chloride, epidermal growth factor (EGF), 0.25% trypsin, bovine pituitary extract (BPE), Greiner Bio-One CELLSTAR µClear™ 96-well, Cell Culture-Treated Flat-bottom Microplates, heat-inactivated fetal bovine serum (FBS), CellMask Orange, 4′, 6-diamidino-2-phenylindole (DAPI), Gibco™ horse serum, bicinchoninic acid assays (BCA), Antibiotic-Antimycotic (100X), Basix Syringe Filters (PVDF; 0.45 µm, 0.22 µm) MilliporeSigma Steriflip Sterile Disposable Vacuum Filter Units (0.45 µm), B-PER Complete, LIVE/DEAD Fixable Near-IR Dead Cell Stain kit, and Vybrant DiD Cell-Labeling Solution were sourced from ThermoFisher Scientific (Waltham, MA). New York City III (NYCIII) media components (HEPES, proteose peptone, sodium chloride, dextrose, and yeast extract), poly-L-lysine, Paraformaldehyde, carbenicillin disodium salt, Amicon Ultra Centrifugal filters (10 kDa MWCO), and Whatman Puradisc 30 syringe filters (5 µm, 1.2 μm) were obtained from MilliporeSigma (Burlington, MA). Formvar/Carbon 200 Mesh grids were sourced from Electron Microscopy Sciences (Hatfield, PA). 32 mL Open-Top Thickwall Polycarbonate Tubes (25 × 89mm) and 38.5 mL Open-Top Thinwall Ultra-Clear Tubes (25 × 89 mm) were sourced from Beckman Coulter (Indianapolis, IN). Uranyl acetate was provided by the Laboratory for Biological Ultrastructure at the University of Maryland. µ-Slide 18-well glass bottom imaging wells were purchased from Ibidi (Fitchburg, WI). Cell Counting Kit-8 was purchased from APExBio (Houston, TX). Calcium Chloride solution (0.5 M) was purchased from PromoCell (Heidelberg, Germany). Cole-Parmer Essentials Disposable Bottle Top Aspirators (PVDF, 0.22 µm) were purchased from Cole-Parmer (Vernon Hills, IL). XenoLight DiR fluorescent dye was sourced from Revvity (Waltham, MA). QIAprep Spin Miniprep Kits were sourced from Qiagen (Venlo, The Netherlands). TGX Stain-Free™ gels, Clarity Max Western ECL Substrate, and Goat Anti-Rabbit IgG (H+L)-HRP Conjugate antibody were sourced from Bio-Rad (Hercules, CA). LB Miller Broth was sourced from MP Biomedicals (Solon, OH). Isopropyl β-d-1-thiogalactopyranoside (IPTG) was purchased from IBI Scientific (Dubuque, Iowa).

### Plasmid assembly for protein production and periplasmic translocation of moxNeonGreen (moxNG)

Plasmids pHKSM010 and pVHSM035 were used to express moxNG and moxNG-6xHisTag, respectively, each fused to the C-terminus of the OmpA signal peptide (OmpA_SP_) for periplasmic translocation. Plasmid pHKSM010 was assembled by inserting OmpA_SP_-<u>moxNG</u> into the pQE80 backbone. The C-terminal 6xHisTag was added to pHKSM010 to create pVHSM035. Plasmids pRASM009 and pRASM010 were generated by removing OmpA_SP_ from pHKSM010 and pVHSM035, respectively. Plasmid pVHSM045, used to purify moxNG for western blot standard curves, was constructed by replacing sf-GFP with moxNG pSMCAF015.^29^ All the plasmids were constructed using Golden Gate Assembly with BsaI recognition sites and transformed into chemically competent *E. coli* 5-alpha cells (New England Biolabs, C2987H). Individual colonies were grown overnight, and plasmid DNA was extracted using the Qiagen QIAprep Spin Miniprep Kit. The resulting plasmids were verified via Oxford Nanopore Technologies and subsequently transformed into *E. coli* Nissle 1917. Sequences and primers used are supplied in supplementary information (**Supp. Table 1, Supp. Table 2**).

### Bacterial growth

Particle-free LB Miller media was prepared via ultracentrifugation at 107,000 × g for 12-16 h at 4°C and subsequently filtered using a 0.22 μm sterile filter.^30^ Strains were cultured under aerobic conditions at 37°C, with a shaking speed of 250 rpm. Single colonies transformed with plasmids were inoculated into 35 mL of particle-free LB Miller media containing 100 ug/mL carbenicillin to ensure plasmid retention. After the cultures reached an optical density (OD) of 0.4-0.6, cultures were induced with relevant concentrations of IPTG, and the culture was moved to its final growth temperature. bEVs were isolated from conditioned media after 48 h of growth.

### bEV isolation

bEVs were isolated from conditioned supernatants via ultracentrifugation as previously described, with modifications.^28^ Briefly, cultures were centrifuged at 4,000 × g for 15 min to remove whole cells. Supernatants were pelleted again at 4,000 × g for 15 min. Supernatants were sequentially filtered through 5 μm, 1.2 μm, 0.45 μm syringe filter units, 0.45 μm SteriFlip units, and 0.22 μm syringe filter units before ultracentrifugation at 25,000 × g for 40 min to remove aggregates or additional cell debris. Supernatant was then ultracentrifuged at 107,000 × g for 3 h. The bEV pellet was resuspended in PBS and ultracentrifuged again at 107,000 × g for 3 h as a wash. Supernatant was removed and ∼1 mL of pellet was resuspended and moved to a clean Eppendorf LoBind microcentrifuge tube. Final resuspension volumes were noted for normalization to initial concentration in conditioned media. bEVs were stored at 4°C until characterized, and -80°C for long term storage. Samples were pooled for experimental dosing and TEM imaging and characterized within 24 h (n = 3-6).

### bEV characterization

bEVs were characterized via nanoparticle tracking experiments using a ZetaView® Mono Nanoparticle Tracking Analysis (NTA) Microscope (Particle Metrix) equipped with a 488 nm laser. EVs were diluted to a final concentration of 10^7^ particles/mL using 0.1 mM NaOH. Size, concentration, and ζ-potential readings were calculated across 11 cell positions, with sensitivity set to 85, minimum brightness set to 65, minimum area to 10, and maximum area set to 1000.^31^ bEV concentrations were normalized to initial culture volume using the resuspension volume. To evaluate the morphology of isolated bEVs, we utilized transmission electron microscopy (TEM). TEM was completed at the University of Maryland Laboratory for Biological Ultrastructure. Samples were prepared at room temperature by depositing pooled samples on Carbon/Formvar 200 mesh grids. Samples were negatively stained using 1% uranyl acetate. Nanoparticles were confirmed to be bEVs based on images taken of diluted cultures (**Supp.** Figure 1). Images were taken using a Hitachi TEM HT7700 microscope. Finally, we used a bicinchoninic acid (BCA) assay to measure protein content in the isolated bEVs. The BCA assay was completed according to manufacturer instructions. Briefly, 25 μL of sample was incubated with the working solution for 30 min at 37°C. Absorbance measurements were taken at 547 nm in a TECAN SPARK plate reader (n = 6-7).

### moxNG purification

A single colony of *E. coli* BL21(DE3) (New England Biolabs) harboring plasmid pVHSM045 for expression of moxNG-HisTag was inoculated in 500 mL of LB Broth. After ∼16 h of growth at 37°C and a shaking speed of 250 rpm, cells were induced with 0.2% w/v L-arabinose. Cells were harvested after 48 h by centrifugation at 10,000 × g for 5 min, resuspended in lysis buffer (50 mM Tris pH 8.0, 300 mM NaCl, 5% v/v glycerol, and 10 mM Imidazole), and lysed using Fisher Scientific 550 Sonic Dismembrator. The lysate was centrifuged at 12,000 × g for 1 h, and the supernatant was collected for protein purification. The proteins were purified using a Ni Sepharose™ 6 Fast Flow (Cytiva) column and buffers containing 50 mM Tris pH 8.0, 300 mM NaCl, 5% v/v glycerol, and 10-250 mM Imidazole. Protein purity was confirmed by SDS-PAGE.

### Determination of moxNG loading

In addition to the total concentration of bEVs, we measured the population of bEVs loaded with moxNG via NTA. Fluorescent particle populations were activated by the standard 488 nm laser, and moxNG emissions were isolated due to the addition of a 500 nm filter. Again, EVs were diluted to a final concentration of 10^7^ particles/mL using 0.1 mM NaOH. Fluorescent population concentrations, size, ζ-potential readings and were calculated across 11 cell positions, with sensitivity set to 91, minimum brightness set to 20, minimum area to 10, and maximum area set to 1000. Loading percentage was calculated as the fluorescent population of bEVs over the total population of bEVs. All frames were manually accepted to determine the concentration of bEVs, potentially resulting in an overestimation of populations below 3% Loading. Size and ζ-potential are not reported for samples below 20 particles/frame (n = 6-9).

For western blot analysis, cells transformed with pVHSM035 were cultured as stated previously. Culture samples were lysed using B-PER Complete. The protein content of the cell lysate was quantified using BCA analysis and normalized prior to loading. bEV samples after isolation were characterized as mentioned previously and normalized to 1.3×10^10^ bEVs before loading. Purified moxNG was loaded in concentrations 0.5-15 ng/µL to create a standard curve for quantification of moxNG content in cells and bEVs. Gels were run at 120 V, after which they were transferred to 0.2 μm nitrocellulose membrane and blocked at room temperature for 3 h with 1% BSA. Membranes were then washed in TBST buffer four times before incubation in a 1:250 dilution of Rabbit Anti-HisTag overnight at 4°C with gentle agitation. Membranes were washed in TBST buffer four times before incubation in a 1:2000 dilution of Goat Anti-Rabbit IgG (H+L)-HRP Conjugate antibody (BioRad) solution for 1 h at room temperature. Membranes were washed in TBST buffer four times before Clarity Max Western ECL Substrate (BioRad) was applied. The membrane was imaged immediately for chemiluminescence. Three technical replicates were averaged for reporting (n = 6).

### Confocal microscopy

To visualize moxNG within cells, single colonies were used to inoculate 35 mL of LB Miller media. After the cultures reached an optical density of 0.4-0.6, cultures were induced with relevant concentrations of IPTG. The cells were allowed to grow for 3 h before collection. 2 mL of culture was aliquoted and washed three times with 1 mL of 0.01 M PBS. Then, a ∼2 μL was placed between agarose (1.5% w/v) and a glass coverslip. Images were acquired using the Olympus FV3000 confocal microscope and FLUOVIEW software.

### bEV labeling

bEVs were labeled with DiD dye for uptake experiments, and DiR dye for *in vivo* biodistribution. To achieve necessary concentrations, bEVs were concentrated using 10 kDa MWCO filters at 3000 × g. Briefly, bEVs and dye were incubated at 37°C for 20 minutes at a ratio of 1.5 μL dye per 100 μL bEV sample. The sample was washed three times in a 10 kDa MWCO filter with PBS at 3000 × g. Samples were filtered until the last 500 μL remained. Resulting concentrations were determined via NTA. For *in vivo* studies, PBS was processed with the DiR dye in the same manner as bEVs.

### Cell culture

The VK2/E6E7 vaginal epithelial cells were cultured in keratinocyte-serum free medium supplemented with 0.1 ng/mL EGF, 0.05 mg/mL BPE, 44.1 mg/L calcium chloride, and 1% Antibacterial-Antimycotic. Experiments utilized passages 5-15. Similarly, End1/E6E7 endocervical cells were cultured in keratinocyte-serum free medium without calcium chloride, supplemented with 0.1 ng/mL epidermal growth factor, 0.05 mg/mL BPE, 44.1 mg/L calcium chloride, and 1% Antibacterial-Antimycotic. Experiments utilized passages 4-10. All cells were maintained at 37°C and 5% CO^2^, according to manufacturer instructions. Media was exchanged every 48 h. Cells were passaged at 85-95% confluency.

### Cell viability assay

For cytotoxicity assays, cells were seeded into 96-well plates at 0.04 × 10^6^ cells/well. Cells were allowed to adhere for 24 h. Media was exchanged immediately before dosing with 10 μL of bEVs for a final concentration of 2 × 10^8^, 2 × 10^7^, or 2 × 10^6^ bEVs per well, equivalent to 5000, 500, or 50 bEVs/cell. After 24 h incubation with bEVs, cell viability was determined using Cell Counting Kit-8 (CCK-8) according to manufacturer instructions. Briefly, 10 μL of reagent was added to each well and incubated at 37°C for 1 h. Absorbance was read at 450 nm. The percentage of viable cells was determined by comparison to the vehicle control group (PBS) (n = 8).

### bEV uptake quantification

To measure the efficiency of cellular uptake, cells were seeded into 96-well plates at 0.04 × 10^6^ cells/well and allowed to adhere overnight. After 24 h, media was replaced, and cells were dosed with DiD-labeled bEVs at 5000 bEVs/mL. After the allotted time, cells were washed three times with PBS, before being incubated at RT with 1% Triton. Standards were made with known concentrations of particles diluted in 1% Triton. Fluorescence readings for DiD were done in a TECAN SPARK plate reader at an EX:644 nm, EM:688 nm, and bandwidth of 5 nm (n = 8).

Additionally, we used flow cytometry to quantify the number of cells that internalized bEVs. Cells were seeded to 96-well plates at 0.04 × 10^6^ cells/well and allowed to adhere overnight. In the morning, media was replaced, and cells were dosed with DiD-labeled bEVs at 5000 bEVs/mL. At each timepoint, media was removed, and cells were lifted via 0.25% trypsin. The reaction was quenched with DMEM + 10% FBS. Cells were washed once with PBS containing 1% w/v BSA (FACS buffer) and stained with LIVE/DEAD Fixable Viability Dye for 30 minutes at 4°C. Cells were then washed with FACS buffer two times and were analyzed immediately with a CytoFLEX (Beckman Coulter). Flow cytometry data analysis was performed using FlowJo software (Version 10, FlowJo LLC). Three wells were combined per replicate (n=8).

To visualize cellular uptake, glass bottom well plates were treated with poly-L-lysine for 20 min and dried overnight. Cells were seeded at 0.04 × 10^6^ cells/well and allowed to adhere for 24 h. Once adhered, media was exchanged, and cells were dosed at 5000 bEVs/cell. After 24 h, cell membranes were labeled via CellMask Orange membrane stain according to manufacturer’s instructions. Cells were fixed via 2% w/v formaldehyde at 37°C for 10 min, washed with PBS 3x, and incubated in 300 nM DAPI for 5 min. Cells were imaged using a LSM 980 Laser Scanning Confocal microscope (Zeiss, Oberkochen, Germany; n = 3 images per well, 3 wells per condition).

### bEV compatibility with vaginal microbes

NYCIII media (0.4% w/v HEPES, 1.5% w/v proteose peptone, 0.5% w/v sodium chloride, 0.5% w/v dextrose, 2.6% w/v yeast extract) was supplemented with 10% v/v horse serum and prepared according to ATCC instructions. *Lactobacillus crispatus* was plated on NYCIII 1.5% agar plates, and cultured anaerobically (5% Hydrogen, 95% Nitrogen) for 72 h. Colonies were passaged to 5 mL of NYCIII and grown for 72 h until confluency. For compatibility experiments, cultures were diluted to an OD600 of 0.05. 200 μL of diluted culture and dosed with 2×10^8^ bEVs, 2×10^7^ bEVs, 2×10^6^ bEVs, or PBS, in 20 μL.

To measure the colony forming units (CFUs), *L. crispatus* cultures were exposed to bEVs (doses) for 8 h at 37°C in anaerobic conditions. After incubation, 10 μL of diluted culture was deposited on NYCIII agar and allowed to grow for 48 h, at which time CFUs were counted. Dilutions resulting in 4-40 CFUs were averaged to determine CFU/mL (n = 4-8).

To measure the OD, *L. crispatus* cultures were seeded in a 96-well plate and housed in a Cerillo Discovery, at 37°C in anaerobic conditions. OD600 readings were taken every 5 min for 48 h. OD600 readings were normalized to the initial reading (n =n5).

### Biodistribution of bEVs *in vivo*

C57Bl/6 female mice between 12-16 weeks old were used for biodistribution studies. On Day -3, mice were subcutaneously injected with 100 µL of 1 mg/mL estradiol to sync estrus cycles. On Day 0, mice were dosed vaginally with 3.75 × 10^10^ DiR-labeled bEVs. At each timepoint, mice were euthanized and the reproductive tract, including vaginal, cervix, and uterine horn, was removed imaged using an *In Vivo* Imaging System (n = 2 for PBS, n = 4 for bEVs). Animals used in this study were euthanized using a precision vaporizer with an induction chamber and waste gas scavenger in which isoflurane was administered in 2.5% O_2_ for >2 min. Cervical dislocation was performed to ensure euthanasia. All procedures in this study were approved by the University of Maryland Baltimore Institutional Animal Care and Use Committee. All procedures were conducted in accordance with the National Institutes of Health Guide for the Care and Use of Laboratory Animals. Reporting of animal experiments is done in accordance with ARRIVE guidelines.

### Statistical analyses

Standard quantitative analyses were performed using analysis of variance (ANOVA) testing with appropriate repeated measures. Tukey post-hoc tests were used with significance at *p* < 0.05. Unpaired two-tailed T-tests were used when applicable, with significance at *p* < 0.05. A maximum of one outlier in each group was removed using Grubb’s outlier tests with alpha = 0.05. Data are represented as mean ± standard error of the mean.

## Results

### Genetic engineering facilitates bEV loading

Given the spectrum of genetic tools available for *E. coli*, we adopted *E. coli* Nissle 1917 (EcN) as a foundational platform for engineering bEVs into vaginal drug delivery vehicles. Moreover, the EcN strain has beneficial effects on mammalian cells, promoting its wide use as a safe probiotic.^32^ In Gram-negative bacteria such as *E. coli*, bEVs originate from the outer membrane.^33-35^ Consequently, to enable loading of target proteins into bEVs, these proteins must first be translocated into the periplasmic space (**Figure 1A**). To demonstrate the feasibility of this strategy, we engineered the periplasmic expression of moxNeonGreen (moxNG) as a model protein. Periplasmic targeting was achieved by fusing moxNG to the C-terminus of the *E. coli* OmpA signal peptide OmpA_SP_; **Figure 1B**). ^36,37^ Protein expression was placed under the control of an Isopropyl-β-D-thiogalactopyranoside (IPTG)-inducible promoter, allowing modulation of expression levels in subsequent optimization studies. Confocal microscopy revealed that cytosolic expression of moxNG (C) resulted in diffuse fluorescence throughout the cell (**Figure 1C**). In contrast, periplasmic localization of the OmpA_SP_–moxNG fusion protein (P), resulted in fluorescence restricted to the cell periphery (**Figure 1D**, **Supp.** Figure 2). Nanoparticle tracking analysis (NTA) showed an increase in bEV concentration with periplasmic expression of moxNG (1.22 ± 0.30 × 10^10^ bEVs/mL, **Figure 1E**), compared to WT (1.12 ± 0.08 × 10^9^ bEVs/mL, *p* = 0.0022), and cytosolic expression (9.37 ± 1.33 × 10^8^ bEVs/mL, *p* = 0.0019). Additionally, periplasmic expression resulted in a decrease in size (159.5 ± 7.89 nm, **Figure 1F**), compared to WT (192.4 ± 2.60, *p* = 0.0016), and cytosolic expression (207.5 ± 2.90, *p* < 0.0001). NTA confirmed that the cytosolic expression of moxNG resulted in minimal loading (0.52 ± 0.01%; **Figure 1G**), compared to the 5.16 ± 0.58% obtained by the periplasmic expression (*p* < 0.0001).

**Figure 1:**
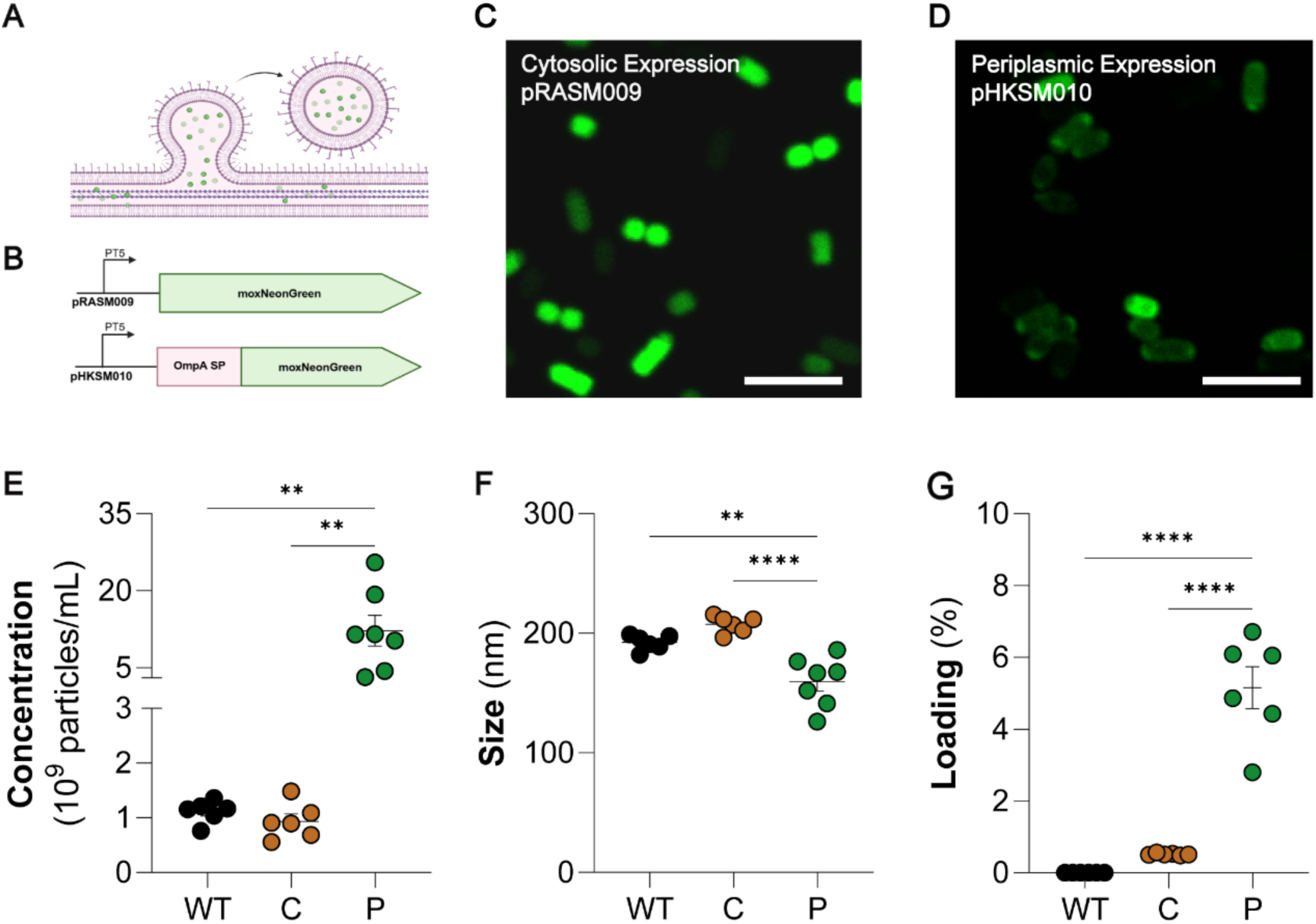
Periplasmic expression of moxNG increases bEV loading. (**A**) Schematic depicting the production of bEVs in Gram-negative bacteria such as *E. coli*, and the loading of periplasmic contents. The outer membrane blebs outward, allowing for the loading of proteins located in the periplasmic space, before pinching off to form a bEV. (**B**) Plasmid constructs can be produced via Golden Gate assembly for the production of moxNG in either the cytosol (pRASM009) or targeted to the periplasmic space via OmpA signal peptide (pHKSM010). Production of moxNG in EcN in the (**C**) cytosol and (**D**) periplasm was confirmed via confocal microscopy. Periplasmic localization is indicated by fluorescence restricted to the cell periphery. Scale bars denote 5 µm. (**E**) Periplasmic expression of moxNG resulted in an increase in bEV concentration (1.22 ± 0.30 × 10^10^), compared to WT (1.12 ± 0.08 × 10^9^, *p* = 0.0022), and cytosolic expression (9.37 ± 1.33 × 10^8^, *p* = 0.0019). (**F**) Periplasmic expression resulted in a decrease in size (159.5 ± 7.89 nm), compared to WT (192.4 ± 2.60, *p* = 0.0016), and cytosolic expression (207.5 ± 2.90, *p* < 0.0001). (**G**) NTA confirmed that the cytosolic expression of moxNG resulted in minimal loading (0.52 ± 0.01%), compared to the 5.16 ± 0.58% obtained by the periplasmic expression (*p* < 0.0001). (n=6-7; \**p* ≤ 0.05, \*\**p* ≤ 0.01, \*\*\**p* ≤ 0.001, and \*\*\*\**p* ≤ 0.0001)

### Optimization of growth conditions for bEV loading

To improve moxNG loading into bEVs, we next optimized expression conditions by varying incubation temperature and IPTG concentration, as high protein production rates are known to saturate the secretion machinery and impair periplasmic translocation.^38-40^ EcN cultures were grown at 18 °C, 25 °C, or 37 °C and induced with low (1.56 nM, denoted as PL), medium (6.25 nM, PM), or high (25 nM, PH) IPTG concentrations.

NTA revealed a consistent inverse correlation between incubation temperature and bEV loading efficiency across all induction levels. Within the PL group, bEVs from cultures grown at 37 °C (PL37, 3.01 ± 0.53%, **Figure 2A**, **Supp.** Figure 3) exhibited lower loading compared to PL25 (15.34 ± 3.30%, *p* = 0.0968) and PL18 (15.74 ± 1.93%, *p* = 0.0989). A similar, statistically significant, trend was observed in the PM group: PM37 bEVs (7.12 ± 2.03%) showed reduced loading compared to PM25 (30.17 ± 5.40%, *p* = 0.0002) and PM18 (39.55 ± 2.89%, *p* < 0.0001). Although the PH group followed the same temperature-dependent pattern, overall loading efficiency decreased relative to PM. Specifically, PH37 bEVs (6.18 ± 0.83%) showed lower loading than PH25 (24.79 ± 7.90%, *p* = 0.0093) and PH18 (25.68 ± 4.04%, *p* = 0.0062). Based on these results, the PM18 condition—6.25 nM IPTG induction at 18 °C—was selected for subsequent experiments, as it yielded the highest bEV loading efficiency.

**Figure 2:**
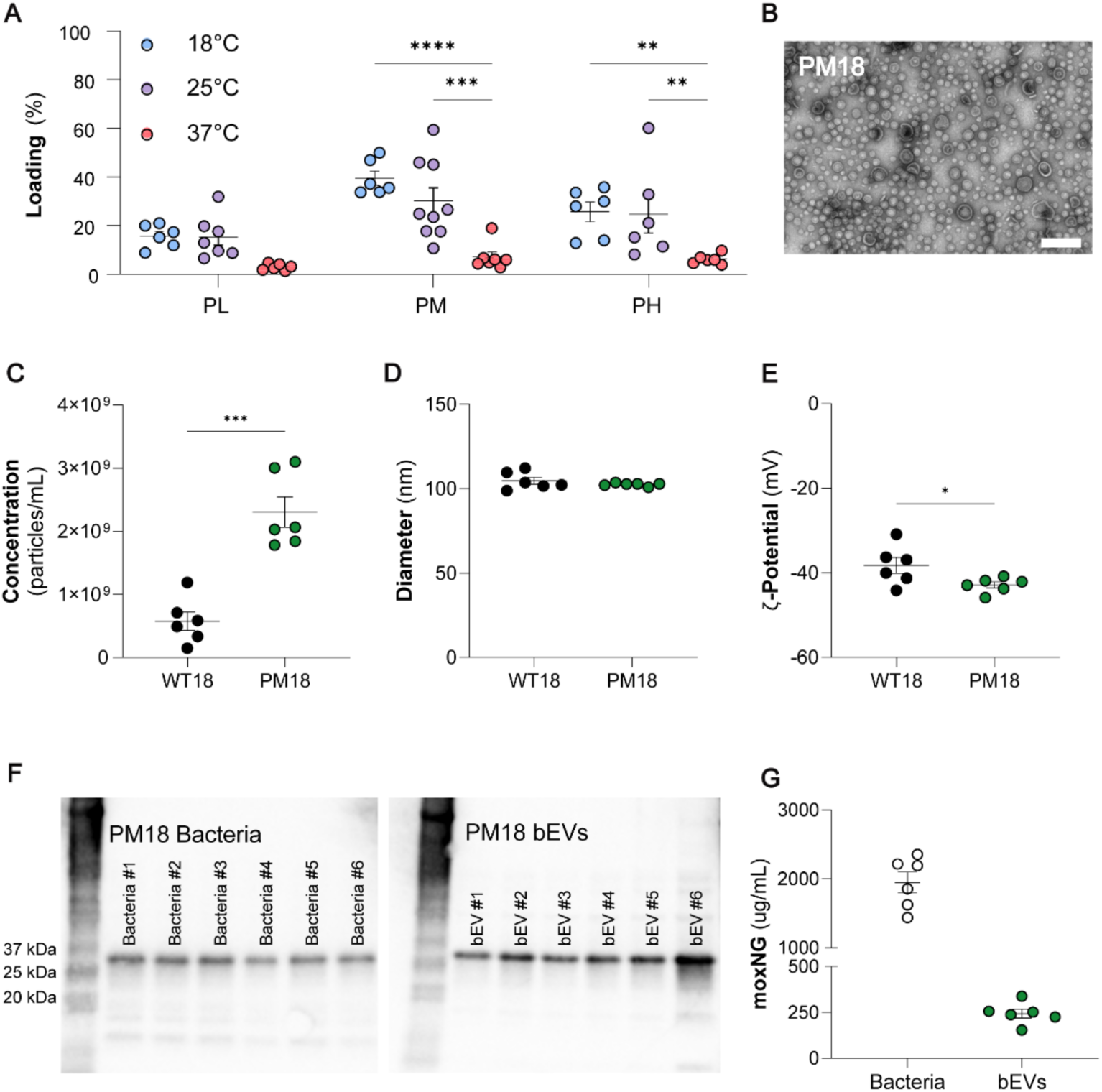
Optimized growth conditions maximize expression of moxNG in bEVs. (**A**) Loading optimization of WT bacteria with periplasmic moxNG expression was determined at 3 different IPTG induction levels, low (1.56 nM; PL), medium (6.25 nM; PM) or high (25 nM; PH), and 3 different temperatures (18°C, 25°C, or 37°C). PM18 bEVs had the highest loading percentage (39.55 ± 2.89%) with the least variability and was used for future experiments. 2-way ANOVA was used with Tukey’s post hoc analysis to determine statistical significance. n = 6-9. (**B**) TEM confirms the presence of membrane bound particles in PM18 samples. Scale bar denotes 200 nm. (**C**) There was no significant difference between the concentration of WT18 bEVs (5.78 ± 1.46 x10^8^ bEVs/mL) and PM18 bEVs (2.31 ± 0.24 × 10^9^ bEVs/mL). (**D**) WT18 bEVs (104.8 ± 2.10 nm) and PM18 bEVs (102.6 ± 0.39 nm) were a similar size. (**E**) No difference in ζ-potential was seen between WT18 (-38.25 ± 1.89 mV) and PM18 (-42.87 ± 0.72 mV) bEVs. (n = 6-7) (**F**) Western blots confirm the presence of the moxNG+6xhistag complex in both PM18 bacteria and isolated PM18 bEVs. Representative images of 3 technical replicates are shown. (**G**) PM18 bacteria were determined to have 1948 ± 149.3 µg/mL, while PM18 bEVs were loaded with 243.7 ± 24.14 µg/mL. Unpaired t-tests were used to determine significance. (\**p* ≤ 0.05, \*\**p* ≤ 0.01, \*\*\**p* ≤ 0.001, and \*\*\*\**p* ≤ 0.0001)

We evaluated the physical characteristics of bEVs produced by WT18 and PM18 cultures, in accordance with guidelines provided by the International Society of Extracellular Vesicles (**Figure 2B-E**).^41^ TEM imaging verified the presence of membrane-bound nanoparticles consistent with previous bEV studies (**Figure 2B**). PM18 bEVs had higher concentrations of bEVs (2.31 ± 0.24 × 10^9^ bEVs/mL), compared to WT18 (5.78 ± 1.46 x10^8^ bEVs/mL, *p* = 0.0001, **Figure 2C**). No difference between WT18 (104.8 ± 2.10 nm, *p* < 0.0001) and PM18 bEVs (102.6 ± 0.39 nm) was seen (*p* = 0.3354, **Figure 2D**). A slight decrease in ζ-potential was seen between WT18 (-38.25 ± 1.89 mV, **Figure 2E**) and PM18 (-42.87 ± 0.72 mV, *p* = 0.0452) bEVs. The addition of a polyhistidine-tag in the periplasmic expression plasmid (moxNG+HisTag) allowed for quantification of moxNG in PM18 bEVs via Western Blot (**Supp.** Figure 4). Western blots confirm the presence of the moxNG+HisTag complex in both PM18 bacteria and isolated PM18 bEVs (**Figure 2F**). PM18 bacteria were determined to have 1948 ± 149.3 µg/mL (**Figure 2G**), while PM18 bEVs were loaded with 243.7 ± 24.14 µg/mL. Based on the determined concentration of moxNG-loaded bEVs, we correlate this to 20.40 ± 2.17 µg/10^10^ moxNG+ bEVs. However, we expected heterogeneous loading in bEVs and assume there is a large portion of bEVs with moxNG below the limit of detection via NTA. Considering the total population of bEVs, this moxNG concentration correlates to 6.48 ± 0.77 µg/10^10^ bEVs.

### EVs are compatible with female reproductive tract cells and key vaginal microbiota

As a vaginal therapeutic, bEVs must be compatible with key reproductive tract tissues, as well as the local vaginal microbiota. Vaginal epithelial cells were dosed with 50, 500, or 5000 bEVs/cell. While there was a significant effect of group (WT18 or PM18, *p* = 0.0066, **Figure 3A**), there was no difference between the growth of the vehicle control (100.00 ± 5.46%) and WT18 groups at 50 bEVs/cell (104.1 ± 5.58, *p* = 0.9286), 500 bEVs/cell (114.9 ± 5.81, *p* = 0.1574), or 5000 bEVs/cell (113.8 ± 4.59, *p* = 0.2135) or the vehicle control and PM18 groups at 50 bEVs/cell (100.9 ± 3.61, *p* = 0.9991), 500 bEVs/cell (98.56 ± 3.95, *p* = 0.9965), or 5000 bEVs/cell (94.44 ± 4.01, *p* = 0.8435). Similarly, endocervical cells were dosed with WT18 and PM18 bEVs (**Figure 3B**). In the WT18 groups, we observed no statistical differences between the growth of the vehicle control (100.00 ± 4.93%) and WT18 dosed at 50 bEVs/cell (114.5 ± 5.69, *p* = 0.5093) and 500 bEVs/cell (132.7 ± 6.49, *p* = 0.1117). Cells dosed with WT18 bEVs at 5000 bEVs/cell resulted in an increase in growth (141.3 ± 7.35, *p* = 0.0012). In the PM18 groups, 50 bEVs/cell (128.1 ± 4.97%, *p* = 0.0445) and 500 bEVs/cell (128.2 ± 7.92%, *p* = 0.0436) led to a growth increase. There was no difference between the vehicle control and PM18 dosed at 5000 bEVs/cell (123.4 ± 10.18%, *p* = 0.1240). Together, these results suggest compatibility with mammalian cells in the vaginal environment.

**Figure 3:**
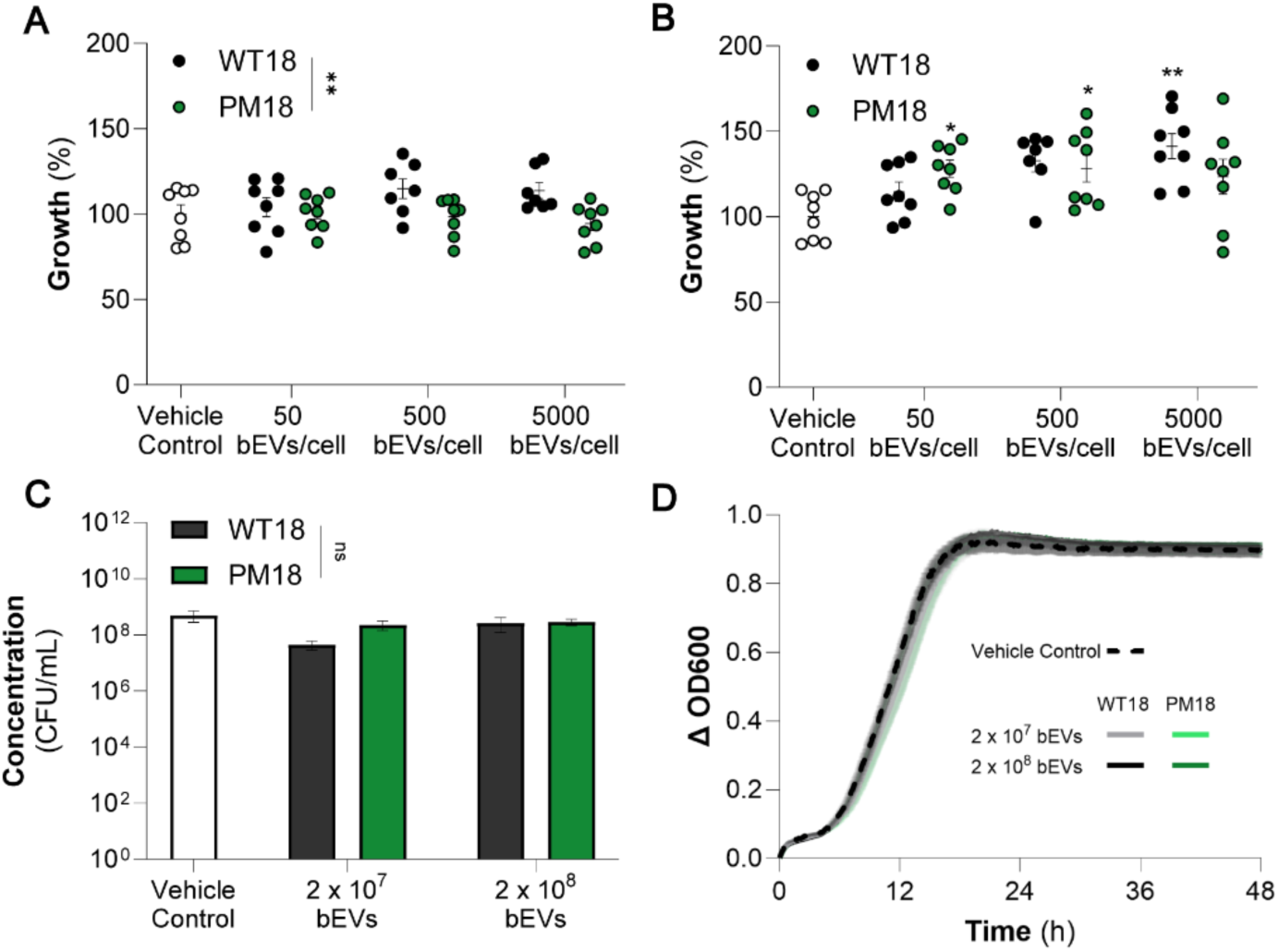
bEVs are compatible with reproductive tract cells and key vaginal microbiota. (**A**) VK2/E6E7 cells were dosed with 50, 500, or 5000 bEVs/cell. No difference was observed between the growth of the vehicle control and WT18 dosed at 50 bEVs/cell, 500 bEVs/cell, or 5000 bEVs/cell (*p* ≥ 0.1574). Similarly, no difference was seen between the vehicle control and PM18 dosed at 50 bEVs/cell, 500 bEVs/cell, or 5000 bEVs/cell *p* ≥ 0.8435). (**B**) End1/E6E7 cells were dosed with WT18 and PM18 bEVs. In the WT18 groups, no difference was seen between the growth of the vehicle control and WT18 dosed at 50 bEVs/cell and 500 bEVs/cell (*p* ≥ 0.1117). WT18 bEVs dosed at 5000 bEVs/cell resulted in an increase in growth (*p* = 0.0012). In the PM18 groups, 50 bEVs/cell and 500 bEVs/cell resulted in a significant increase in growth, compared to the vehicle control (*p* ≤ 0.0445). We observed no difference between the vehicle control and PM18 dosed at 5000 bEVs/cell (*p* = 0.1240). (**C**) No changes in *L. crispatus* CFU concentration were seen at any concentration of WT18 or PM18 bEVs (*p* ≥ 0.5313). (**D**) No differences were seen in the optical density of any group over the course of 48 h. 2-way ANOVA was used with Tukey’s post hoc analysis to determine statistical significance. (\**p* ≤ 0.05, \*\**p* ≤ 0.01, \*\*\**p* ≤ 0.001, and \*\*\*\**p* ≤ 0.0001) (\**p* ≤ 0.05, \*\**p* ≤ 0.01, \*\*\**p* ≤ 0.001, and \*\*\*\**p* ≤ 0.0001)

To investigate the response of bEVs on the vaginal microbiome, we utilized *L. crispatus*, a common bacteria found in a healthy vaginal environment. No difference was seen between the bacterial concentration of the vehicle control group (4.91 ± 2.12 × 10^8^ CFU/mL) and WT18 groups dosed with 2 × 10^7^ bEVs (4.40 ± 1.57 × 10^8^ CFU/mL, *p* = 0.5313) or 2 × 10^8^ bEVs (2.67 ± 1.47 × 10^8^ CFU/mL, *p* = 0.8862). Additionally, no difference was seen in the bacterial concentration of the vehicle control and PM18 dosed at 2 × 10^7^ bEVs (2.28 ± 0.86 × 10^8^ CFU/mL, *p* = 0.6705) or 2 × 10^8^ bEVs (6.73 ± 1.53 × 10^8^ CFU/mL, *p* = 0.8862, **Figure 3C**). No differences were observed in the optical density between groups for 48 h of growth (**Figure 3D**).

### bEVs are internalized by vaginal epithelial cells *in vitro*

As a drug delivery platform, it is critical that bEVs can enter cells to deliver therapeutic cargoes. Here, we investigated the uptake efficiency of vaginal epithelial cells based on a dose of 5000 bEVs/cell (**Figure 4A**). Interestingly, we observed an increase in uptake of WT18 bEVs (6.55 ± 0.32%) compared to PM18 bEVs (0.64 ± 0.14%, *p* = 0.0029) at 6 h. However, by 24 h, the uptake of PM18 bEVs (23.27 ± 3.08%) significantly increased compared to WT18 bEVs (17.91 ± 0.57%, *p* = 0.0078). Flow cytometry was used to determine the percentage of bEV^+^ cells (**Figure 4B**, **Supp.** Figure 5). At the 2 h timepoint, there was an increased percentage of bEV^+^ cells in the PM18 group (39.24 ± 1.23%) compared to WT18 group (3.82 ± 0.31%, *p* < 0.0001). This trend continued for the 6 h (88.80 ± 0.43% v. 51.80 ± 1.64%, *p* < 0.0001) and 12 h timepoints (97.79 ± 0.40% v. 94.20 ± 0.44%, *p =* 0.0074). By 24 h, both treatment groups reached >99% bEV^+^ cells. The internalization of bEVs by vaginal epithelial cells was confirmed via confocal microscopy (**Figure 4C-D**).

**Figure 4:**
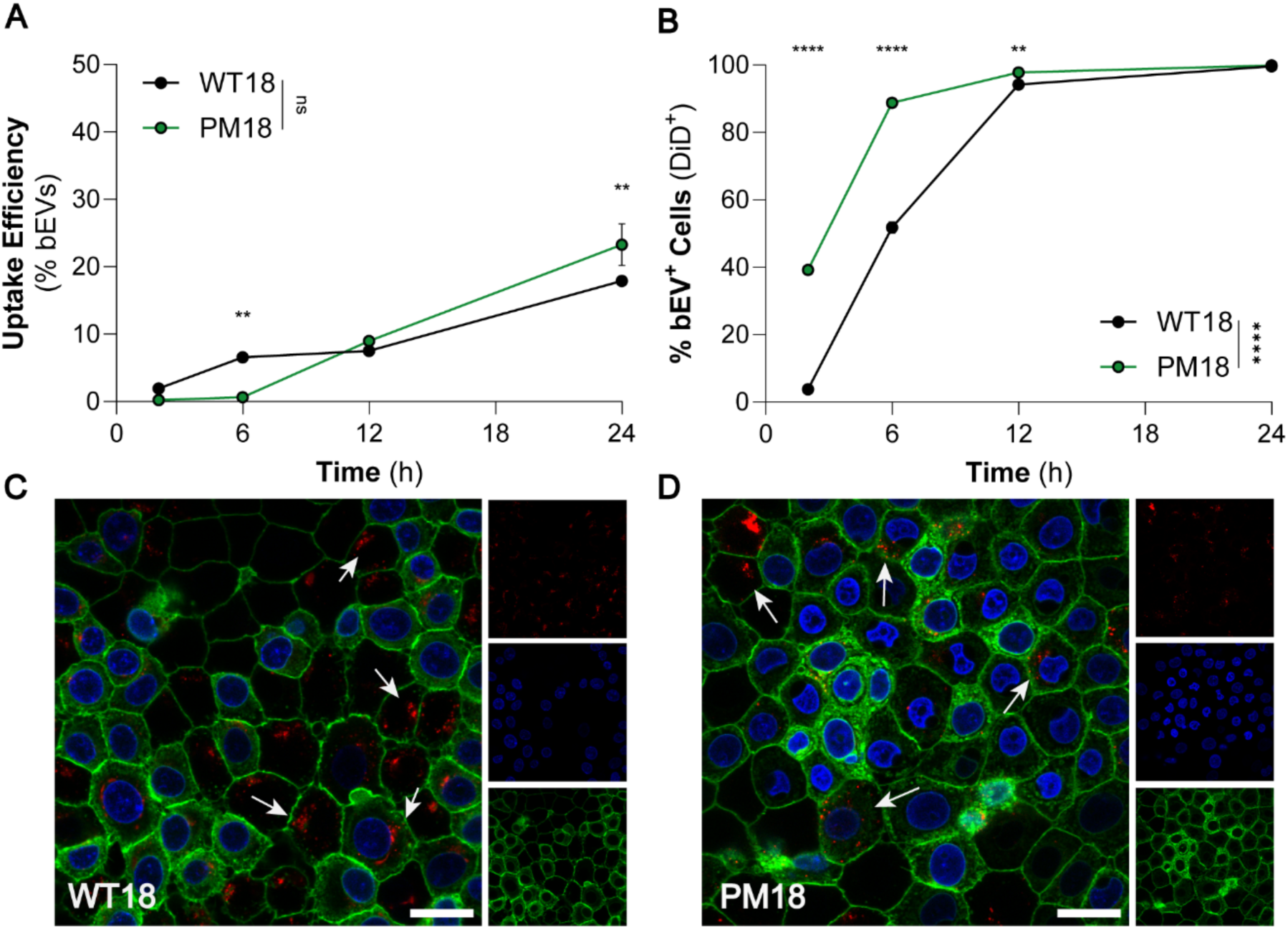
bEVs are internalized by vaginal epithelial cells *in vitro*. (**A**) Uptake efficiency was determined based on a dosage of 5000 bEVs/cell. A lower percentage of PM18 bEVs were internalized at 6 h compared to WT18 bEVs (6.55 ± 0.32% vs 0.64 ± 0.14%, *p* = 0.0029). At 24 h, more PM18 bEVs were internalized compared to WT18 bEVs (17.91 ± 0.57% vs 23.27 ± 3.08%, *p* = 0.0078). (**B**) Flow cytometry was used to determine the percentage of bEV^+^ cells. At 2 h after dosing more cells in the PM18 group (39.24 ± 1.23%) were bEV^+^, compared to the WT18 group (3.82 ± 0.31%, *p* < 0.0001). At 6 h, the PM18 group had more bEV^+^ cells than the WT18 group (88.80 ± 0.43% v. 51.80 ± 1.64%, *p* < 0.0001). At 12 h, 97.79 ± 0.40% of cells in the PM18 group were bEV^+^, compared to 94.20 ± 0.44% of cells in the WT18 group (*p =* 0.0074). By 24 h, both groups reached >99% bEV^+^ cells. The presence of (**C**) WT18 bEVs and (**D**) PM18 bEVs in vaginal epithelial cells at 24 h was confirmed via confocal microscopy. Cell membranes are shown in green (CellMask) and localized around nuclei in blue (DAPI). bEVs are shown in red (DiD). Single channels are provided (red, DiD; blue, DAPI; green, CellMask) Scale bars denote 20 μm. 2-way ANOVA was used with Tukey’s post hoc analysis to determine statistical significance. (\**p* ≤ 0.05, \*\**p* ≤ 0.01, \*\*\**p* ≤ 0.001, and \*\*\*\**p* ≤ 0.0001)

### bEVs are internalized by endocervical cells *in vitro*

Beyond the vaginal epithelium, we sought to evaluate the uptake of bEVs by endocervical cells. Uptake efficiency was determined based on a dose of 5000 bEVs/cell. We observed a significant effect of group (WT18 or PM18, *p* < 0.0001, **Figure 5A**) on the uptake of bEVs over time. PM18 bEVs (18.81 ± 0.98%) were taken up to a greater extent than WT18 bEVs (12.60 ± 0.57%, *p* < 0.0001) at 24 h. As with the vaginal epithelial cells, flow cytometry was used to determine the percentage of bEV^+^ cells (**Figure 5B**, **Supp.** Figure 6). At the 2 h timepoint, a larger percentage of the WT18 group was bEV^+^ (62.53 ± 1.51%) compared to PM18 bEVs (32.20 ± 1.56% bEV^+^ cells, *p* < 0.0001). This trend followed for the 6 h (94.81 ± 0.34% v. 69.43 ± 0.58%, *p* < 0.0001), 12 h timepoints (99.70 ± 0.00% v. 89.98 ± 0.50%, *p =* 0.0074), and 24 h timepoints (99.69 ± 0.11% v. 96.20 ± 0.52%, *p =* 0.0074, **Figure 5B**). The presence of bEVs in endocervical cells was confirmed via confocal microscopy (**Figure 5D**).

**Figure 5:**
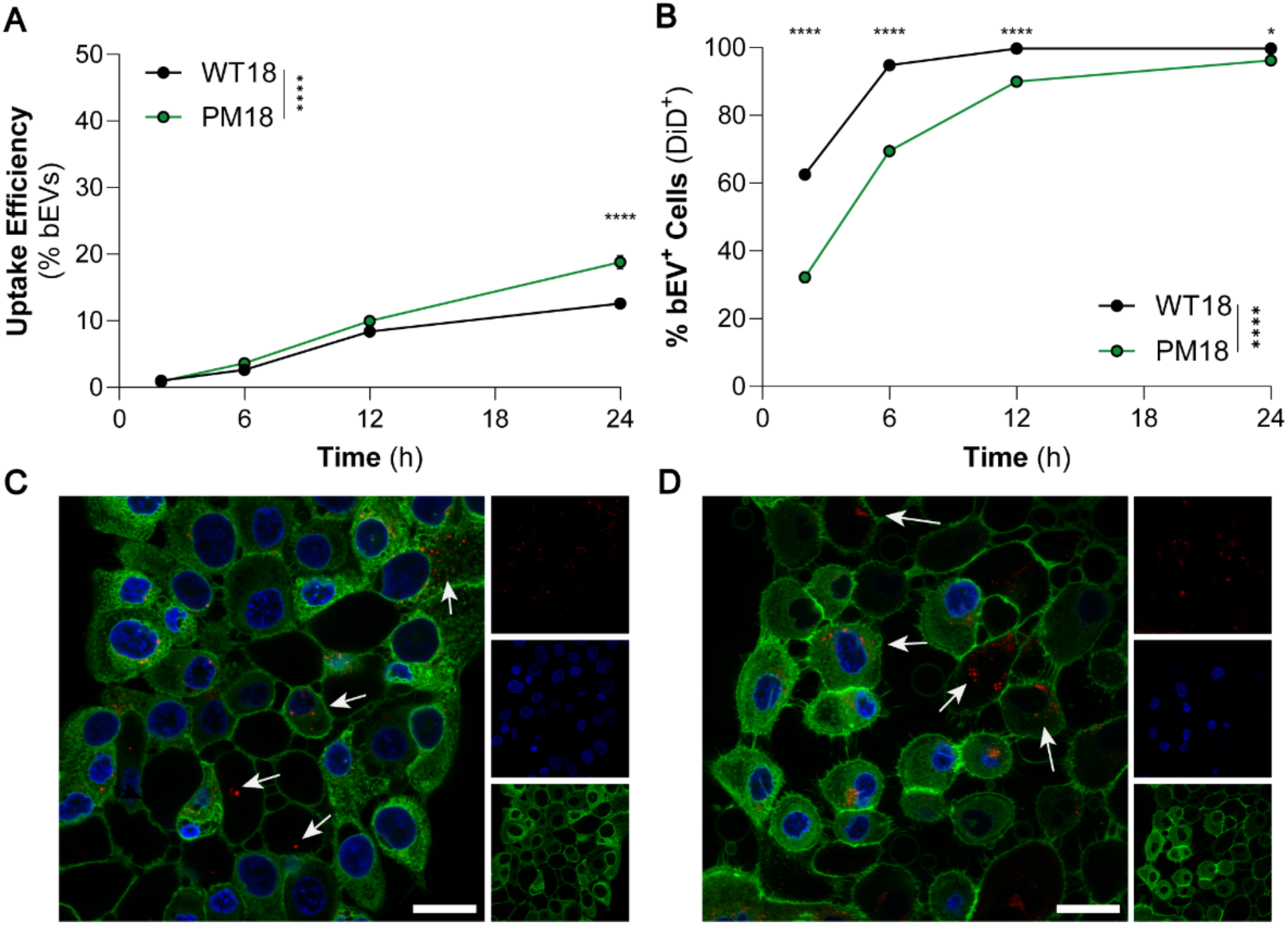
bEVs are taken up by endocervical cells *in vitro*. (**A**) Uptake efficiency was determined based on a dose of 5000 bEVs/cell. PM18 bEVs saw greater uptake compared to WT18 bEVs at 24 h (12.60 ± 0.57% vs 18.81 ± 0.98%, *p* < 0.0001). (**B**) Flow cytometry was used to determine the percentage of bEV^+^ cells. At the 2 h timepoint, WT18 bEVs showed a greater uptake (62.53 ± 1.51% bEV^+^ cells) compared to PM18 bEVs (32.20 ± 1.56% bEV^+^ cells, *p* < 0.0001). This trend followed for the 6 h (94.81 ± 0.34% vs 69.43 ± 0.58% bEV^+^ cells, *p* < 0.0001), 12 h timepoints (99.70 ± 0.00% vs 89.98 ± 0.50% bEV^+^ cells, *p =* 0.0074), and 24 h timepoints (99.69 ± 0.11% vs 96.20 ± 0.52% bEV^+^ cells, *p =* 0.0074) (**E**) Representative flow plots are shown, compared to untreated cells. The presence of (**C**) WT18 bEVs and (**D**) PM18 bEVs in endocervical cells at 24 h was confirmed via confocal microscopy. Single channels are provided (red, DiD; blue, DAPI; green, CellMask). Scale bars denote 20 μm. 2-way ANOVA was used with Tukey’s post hoc analysis to determine statistical significance. (\**p* ≤ 0.05, \*\**p* ≤ 0.01, \*\*\**p* ≤ 0.001, and \*\*\*\**p* ≤ 0.0001)

### bEVs are retained in the vaginal environment *in vivo*

We next sought to evaluate the retention of bEVs in the female reproductive tract *in vivo*. Female mice were dosed vaginally with 3.75 × 10^12^ labeled bEVs, and the female reproductive tract was imaged for bEVs over the course of 24 h. Representative images of vehicle controls (**Figure 6A**) and PM18 groups (**Figure 6B**) are provided. At 2 h, the total radiant efficiency of the vaginal canal for the PM18 group (1.16 ± 0.38 × 10^9^ (p/s)/(µW/cm^2^)) was not significantly different compared to the vehicle control (0.69 ± 0.02 × 10^9^ (p/s)/(µW/cm^2^), *p* = 0.4292, **Figure 6C**). At 6 h, the PM18 group had an increased radiant efficiency (2.04 ± 0.53 × 10^9^ (p/s)/(µW/cm^2^)) compared to vehicle control group (0.65 ± 0.004 × 10^9^ (p/s)/(µW/cm^2^), *p* = 0.0318). By 24 h, the radiant efficiency of the PM18 group decreased (0.90 ± 0.09 × 10^9^ (p/s)/(µW/cm^2^), and was not statistically significant, compared to the vehicle control group (0.67 ± 0.001 × 10^9^ (p/s)/(µW/cm^2^), *p* = 0.6845).

**Figure 6:**
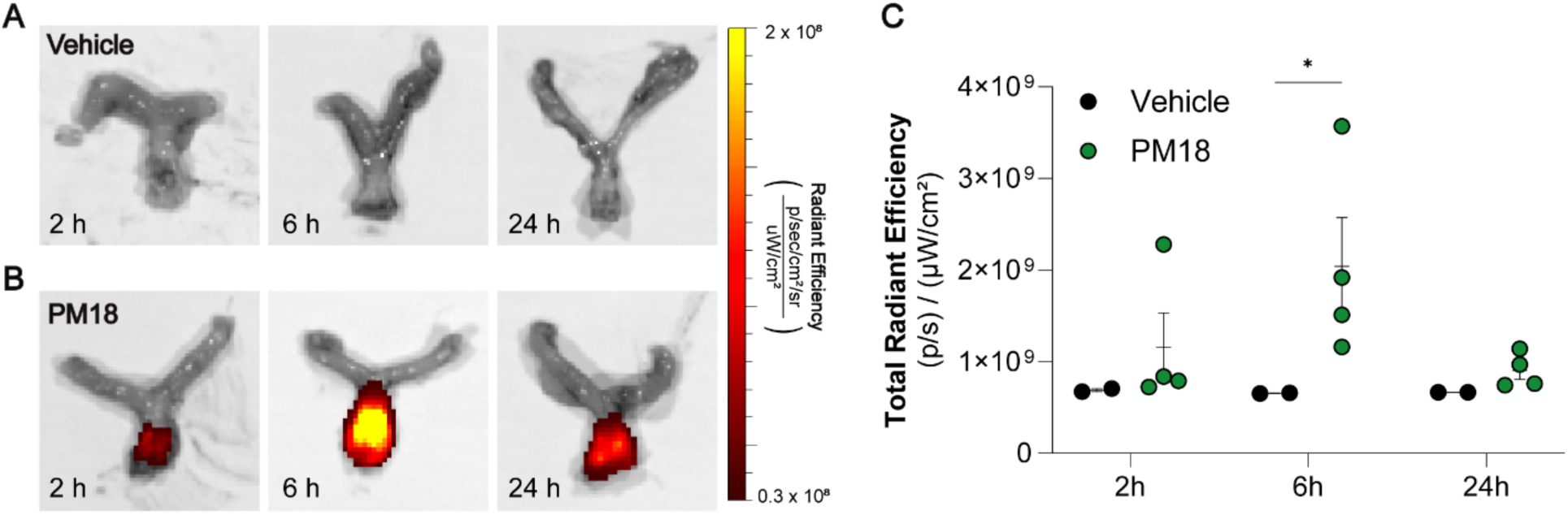
PM18 bEVs are retained in the vagina for 6 h after administration. Representative IVIS images of (**A**) vehicle control (PBS + DiR) and (**B**) PM18 treatment groups. (**C**) At 6 h, the PM18 group exhibited increased radiant efficiency (2.04 ± 0.53 × 10^9^ (p/s)/(µW/cm^2^)) compared to the vehicle control group (0.65 ± 0.004 × 10^9^ (p/s)/(µW/cm^2^), *p* = 0.0318). At 24 h, the radiant efficiency of the PM18 group (0.90 ± 0.09 × 10^9^ (p/s)/(µW/cm^2^) was not statistically different than the vehicle control group (0.67 ± 0.001 × 10^9^ (p/s)/(µW/cm^2^), *p* = 0.6845). 2-way ANOVA was used with Tukey’s post hoc analysis to determine statistical significance. (\**p* ≤ 0.05, \*\**p* ≤ 0.01, \*\*\**p* ≤ 0.001, and \*\*\*\**p* ≤ 0.0001)

## Discussion

This work seeks to establish bEVs as a locally administered nanoparticle platform for treating gynecologic and obstetric diseases. Reproductive conditions are common in women, with 13% of women having fertility issues, 5-20% of women having polycystic ovarian syndrome, and >20% of women having bacterial vaginosis. ^42-44^ It is estimated that endometriosis alone resulted in a loss of 22 billion dollars in 2002.^44,45^ Despite a significant need, there have been few advances in the clinical implementation of next-generation therapies for female reproductive tract diseases.^46^ Preclinical work from our group and others, developed novel nanoparticle formulations for use in treating female reproductive diseases.^9,11,16^ However, translation of these therapeutics is limited by the time and cost requirements associated with the manufacturing workflow. We propose that bEVs may overcome these limitations. Our work here leverages genetic engineering approaches to develop model protein-loaded bEVs, and demonstrates that these bEVs are compatible with the vaginal microenvironment. As there has been no prior report using bEVs to treat female reproductive tract disorders, our work establishes a framework for using bEVs as vaginally delivered therapeutics and sets a foundation for rapidly advancing treatment options for female reproductive tract disorders.

In contrast to synthetic nanoparticles, bEVs represent a nanoparticle platform with endogenous targeting and barrier-crossing abilities, enhanced production capabilities, and reduced associated costs. With industrial biomanufacturing infrastructure in place to produce biopharmaceutics, biochemicals, and food additives, bEVs have a distinct advantage over mammalian EVs.^47^ Compared to cell-based therapies, bEVs exhibit lower immunogenicity and increased stability.^18,48,49^ bEVs have been used in preclinical studies to deliver a range of therapeutic cargoes and have shown promise as a vaccine platform.^17,26,27^ While these studies demonstrate efficacy across several disease indications, the workflow for loading and validating bEV cargoes may hinder clinical translation of bEV-based therapeutics. As we demonstrate in this study, bacteria are amenable to genetic engineering, allowing for cargo loading that is reflected in the resulting bEVs. Here, we use moxNG as a model protein, demonstrating 39.55 ± 2.89% of bEVs grown in optimized conditions carry the target protein. Determination of loading via fluorescent NTA allows for direct quantitative determination of loading, as opposed to semi-qualitative measurements such as western blot. Additionally, fluorescence readings on the NTA allow for the analysis of physiological differences between the total population and fluorescent population, potentially probing for differences in size or ζ-potential that occur as a result of protein loading. We do recognize the limitations of using NTA for loading quantification, as the concentration of moxNG loaded into an individual bEV must reach the fluorescent limit of detection, potentially resulting in underestimation of percent loading. Regardless, our genetic engineering and characterization approaches for protein loading are promising. In particular, genetic engineering strategies, compared to post-isolation modifications, may allow for a drastic reduction of manufacturing costs, enabling more rapid translation of bEV-based therapies.

Vaginal drug delivery is optimal for targeting the female reproductive tract.^50,51^ In contrast to oral or systemic routes of administration, vaginal drug delivery bypasses the acidic and degrading gastrointestinal environment and the hepatic first pass.^50^ Instead, vaginally delivered therapies benefit from the uterine first-pass effect that preferentially circulates vaginally administered drugs to female reproductive tract tissues prior to reaching systemic circulation.^52^ This prevents both the dilution of active pharmaceutical ingredients in the bloodstream, as well as off-target side effects.^50,51^ However, vaginally administered nanoparticles must be compatible with the vaginal microenvironment, including both mammalian and bacterial cells.^53-56^ Our work uses bEVs derived from genetically modified EcN. As other studies have reported vaginal epithelial cell death in response to bEV exposure^57,58^, we thought it important to evaluate the compatibility of EcN-derived bEVs with the vaginal environment. In our work, we observe no detriment to cell viability after administration of both native WT18 bEVs and PM18 bEVs. In addition to biocompatibility with mammalian cells, we demonstrate compatibility with *L. crispatus* cultures. No effect on CFUs is observed after 8 h incubation with WT18 or PM18 bEVs. Additionally, no variation in OD600 is seen over a 48 h time frame. These results, in addition to previous work, underscore the importance of parent cell selection in designing therapeutic bEVs for vaginal drug delivery.

Beyond compatibility with the vaginal microenvironment, we demonstrate that bEVs could serve as a drug delivery platform. The bEVs here were 90-250 nm in size across multiple conditions, consistent with previous reports.^59-61^ This size facilitates increased uptake into female reproductive tract cells.^11,62,63^ Indeed, we observed that bEVs were internalized by over 97% of vaginal epithelial and endocervical cells within 24 h, while causing no cell death. Additionally, we used a preclinical animal model to evaluate the retention of bEVs in the female reproductive tract. bEVs were localized to the vaginal environment and retained for over 6 h. We acknowledge the low power of this study based on a limited sample size. However, at the 2 h timepoint *in vivo* data are consistent with *in vitro* uptake results, and the low signal may be attributed to a low internalization. At 24 h, mice vaginally dosed with DiR-labeled bEVs trended towards an increased signal compared to vehicle controls. Future work may examine differences in membrane composition (lipids, other proteins) that contribute to differences in the internalization of bEVs based on cargo loading (WT18 v. PM18) and cell target (vaginal epithelial v. endocervical). In summary, these data demonstrate the potential for bEVs to deliver therapeutic peptides or biological cargoes to the female reproductive tract.

This work builds a foundation for using bEVs as a vaginal drug delivery platform. Using a bacterial strain with well-defined synthetic biology tools, we demonstrate the use of genetic engineering strategies to yield target protein-loaded bEVs. We show that these bEVs are compatible with the vaginal microenvironment and are retained after vaginal administration to mice. By eliminating the need for post-isolation bioengineering of bEVs, our work has the potential to expedite the translation of treatments for female reproductive tract indications through rapid production of nanomedicines with therapeutic cargoes that can target female reproductive tract cells.

## CRediT Authorship Contribution Statement

**Darby Steinman:** Methodology, Validation, Formal Analysis, Investigation, Writing-Original Draft, Visualization. **Varunaa Sri Hemanth Kumar:** Methodology, Validation, Formal Analysis, Investigation, Writing-Original Draft. **Ryan A. McIlvaine:** Methodology, Validation, Formal Analysis, Writing – Original Draft. **Pranshu Tyagi:** Methodology, Validation, Formal Analysis Investigation, Writing-Review & Editing. **Hahnbit Kang:** Methodology, Validation, Formal Analysis Investigation, Writing-Review & Editing. **Raifah Alam:** Methodology, Validation, Formal Analysis Investigation, Writing-Review & Editing. **Anguo Liu:** Methodology, Validation, Formal Analysis Investigation, Writing-Review & Editing. **Christopher Jewell:** Methodology, Resources, Writing-Review & Editing, Funding Acquisition. **Irina Burd:** Methodology, Resources, Writing-Review & Editing, Funding Acquisition. **Sara Molinari:** Conceptualization, Methodology, Resources, Writing-Review & Editing, Project Administration, Funding Acquisition. **Hannah C Zierden:** Conceptualization, Methodology, Software, Resources, Writing-Review & Editing, Project Administration, Funding Acquisition.

## Acknowledgements

This work was funded by a Faculty-Student Research Award from the Graduate School, University of Maryland (HCZ), the Minta Martin Foundation (HCZ) We acknowledge the support of the University of Maryland, Baltimore, Institute for Clinical & Translational Research (ICTR) and the National Center for Advancing Translational Sciences (NCATS) Clinical Translational Science Award (CTSA), UM1TR004926, as well as the University of Maryland Strategic Partnership: MPowering the State (MPower) (DS, HCZ). DS was supported by the Microbiome Summer Support Fellowship through the Center of Excellence in Microbiome Sciences. Purchase of the Zeiss LSM 980 Airyscan 2 was supported by Award Number 1S10OD025223-01A1 from the National Institute of Health. Funders played no role in the study design, data collection, analysis and interpretation of data, or the writing of the manuscript.

## Data Availability

The data reported here are available upon reasonable request to the corresponding author.

## Supplementary Material

Supporting information is available online.

## Supplementary Material

**Table 1:**
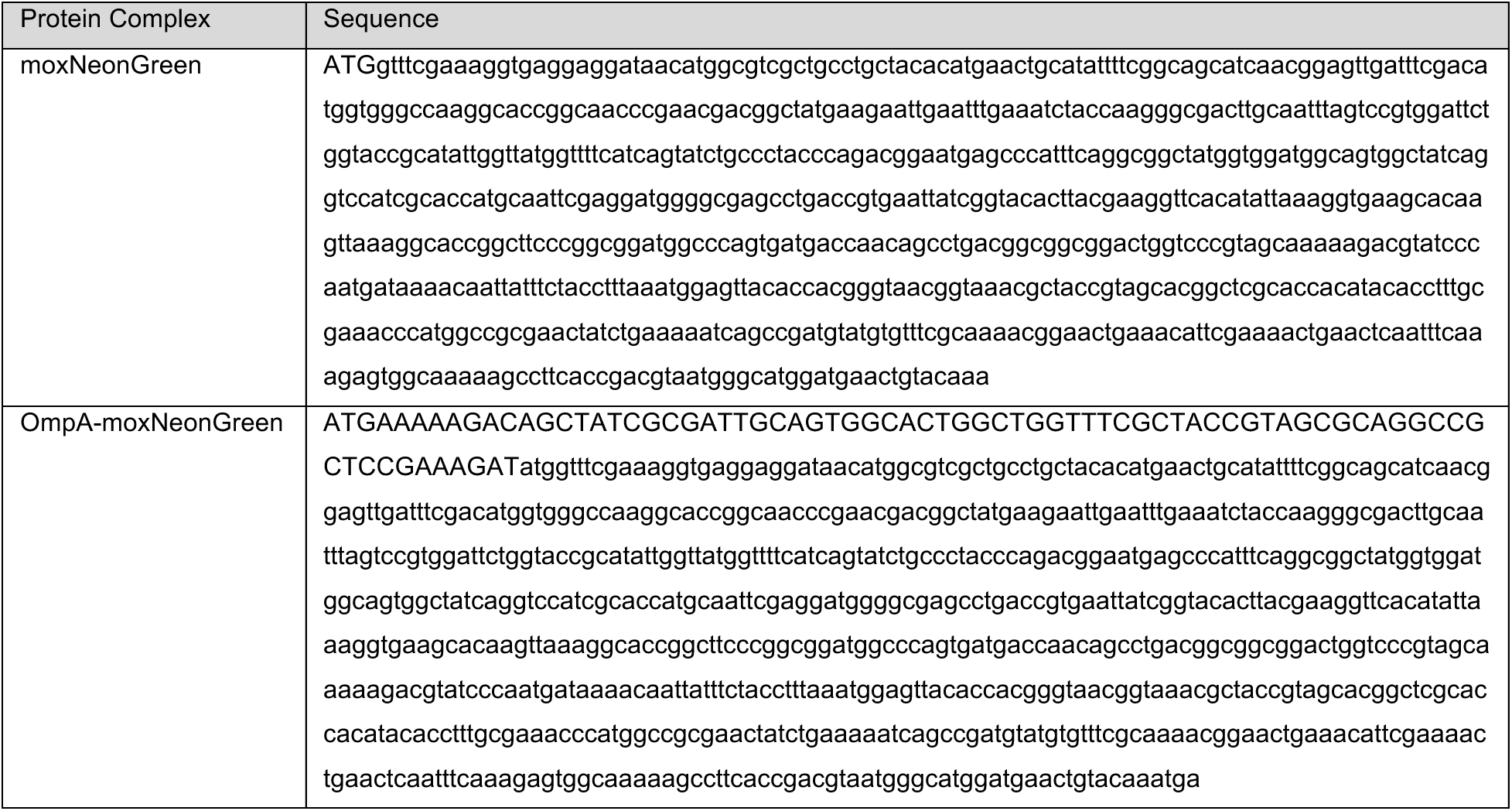
Sequences for relevant proteins.

**Table 2:**
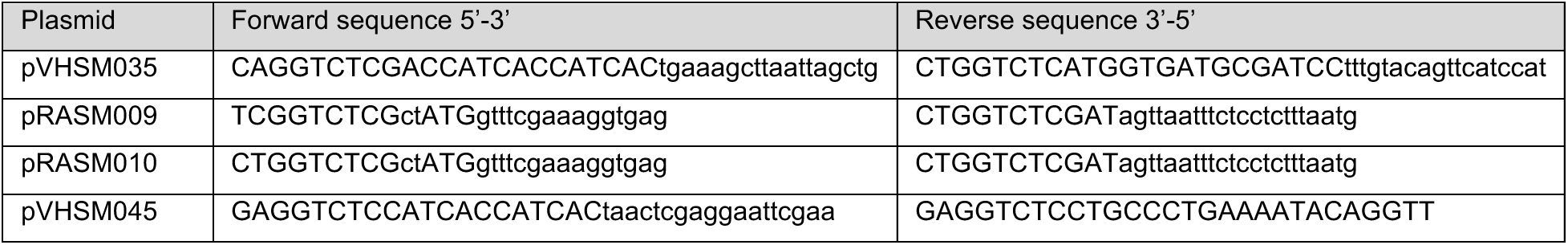
Primers for relevant plasmids.

**Supplemental Figure 1:**
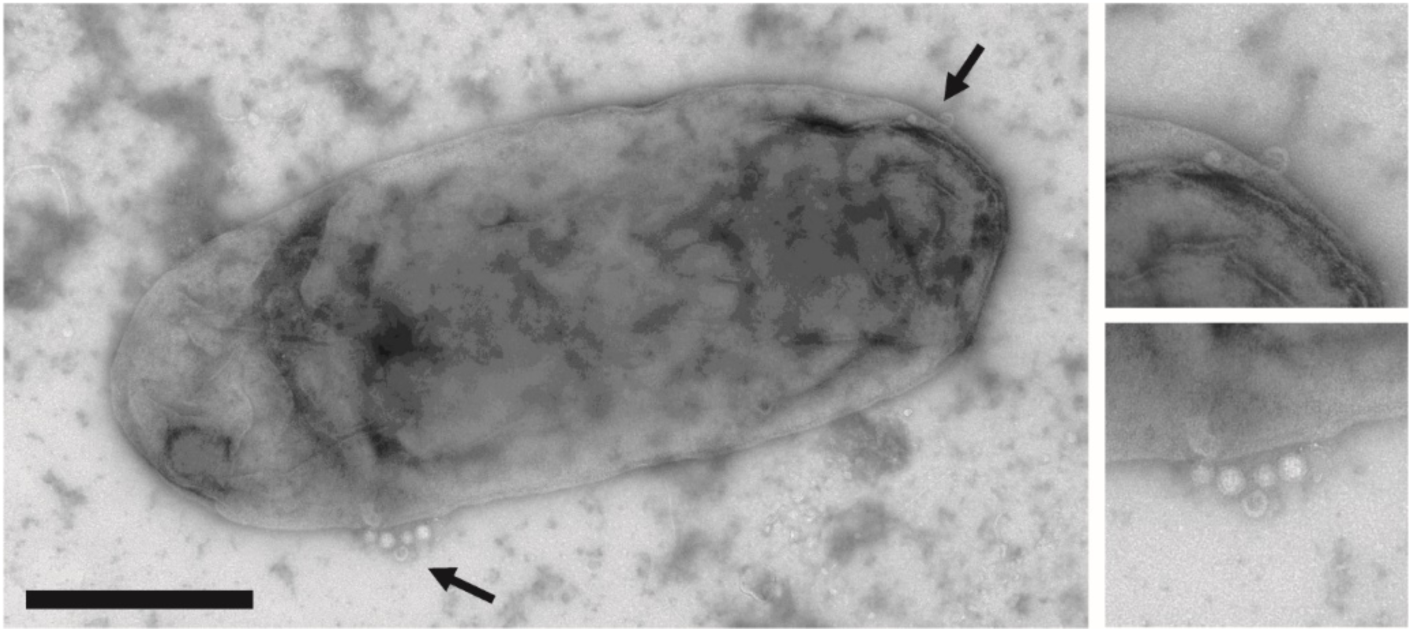
Verification of bEV production by *Escherichia coli* Nissle 1917. Diluted cultures were negatively stained via uranyl acetate and imaged via transmission electron microscopy at the University of Maryland, Laboratory for Biological Ultrastructure. Arrows denote regions of nanoparticles determined to be bEVs. Scale bar denotes 500 nm.

**Supplemental Figure 2:**
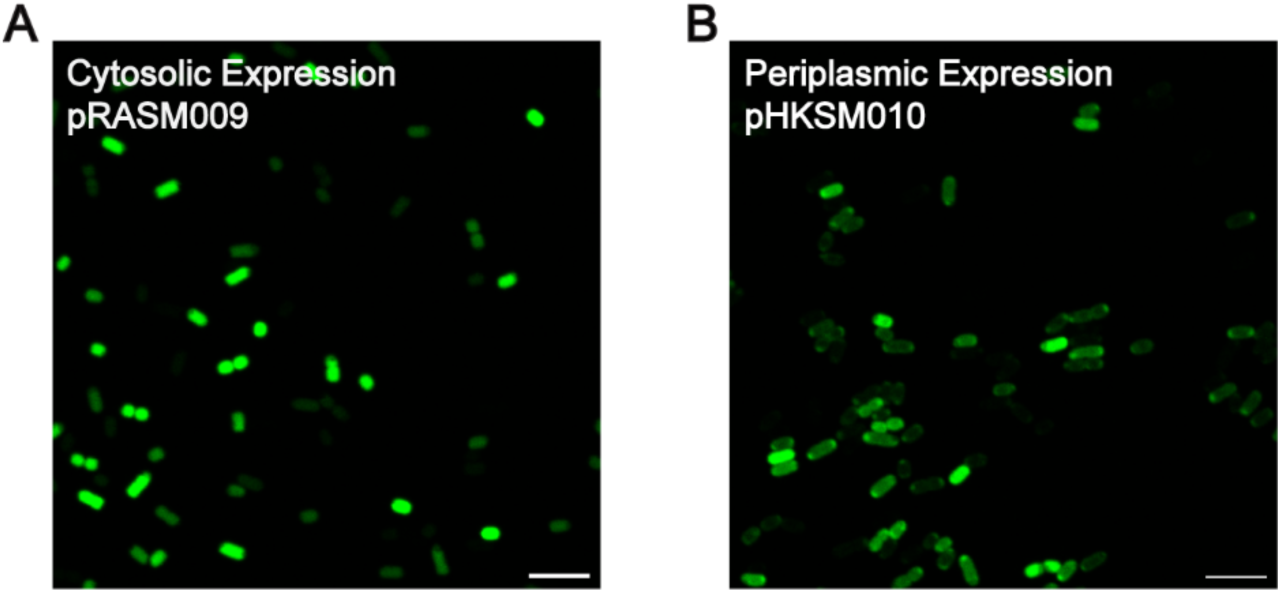
Confocal microscopy confirms moxNG production. Production of moxNG in EcN in the (**A**) cytosol and (**A**) periplasmic was confirmed via confocal microscopy. Periplasmic localization is indicated by fluorescence restricted to the cell periphery. Scale bars denote 5 µm.

**Supplemental Figure 3:**
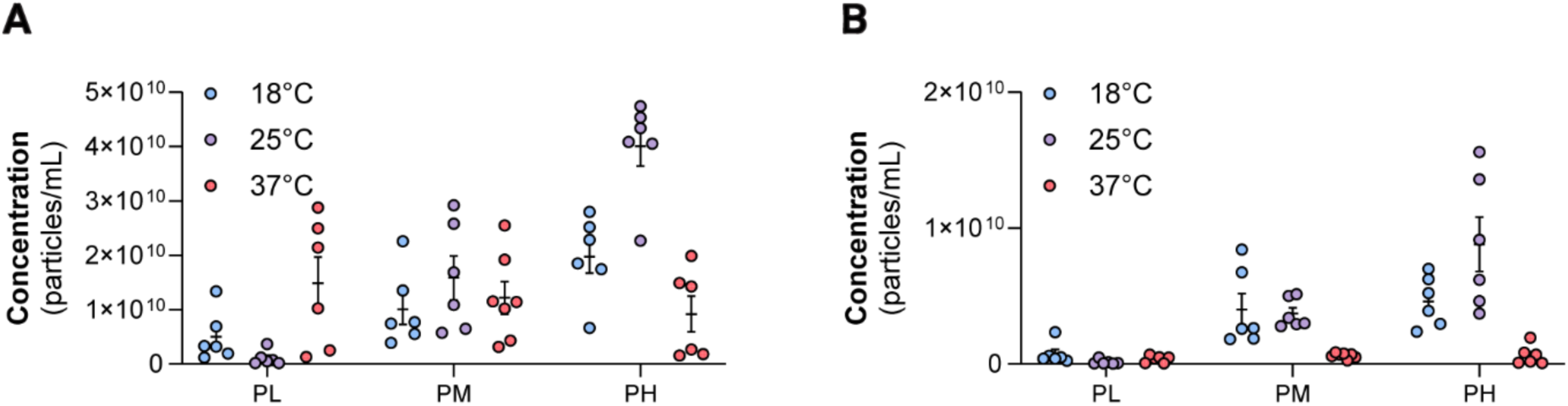
Original bEV concentration for loading optimization experiments. Loading optimization of WT bacteria with periplasmic moxNG expression was determined at 3 different IPTG induction levels, low (1.56 nM; PL), medium (6.25 nM; PM) or high (25 nM; PH), and 3 different temperatures (18°C, 25°C, or 37°C). The concentration of the (**A**) total population and (**B**) fluorescent population were determined via nanoparticle tracking analysis.

**Supplemental Figure 4:**
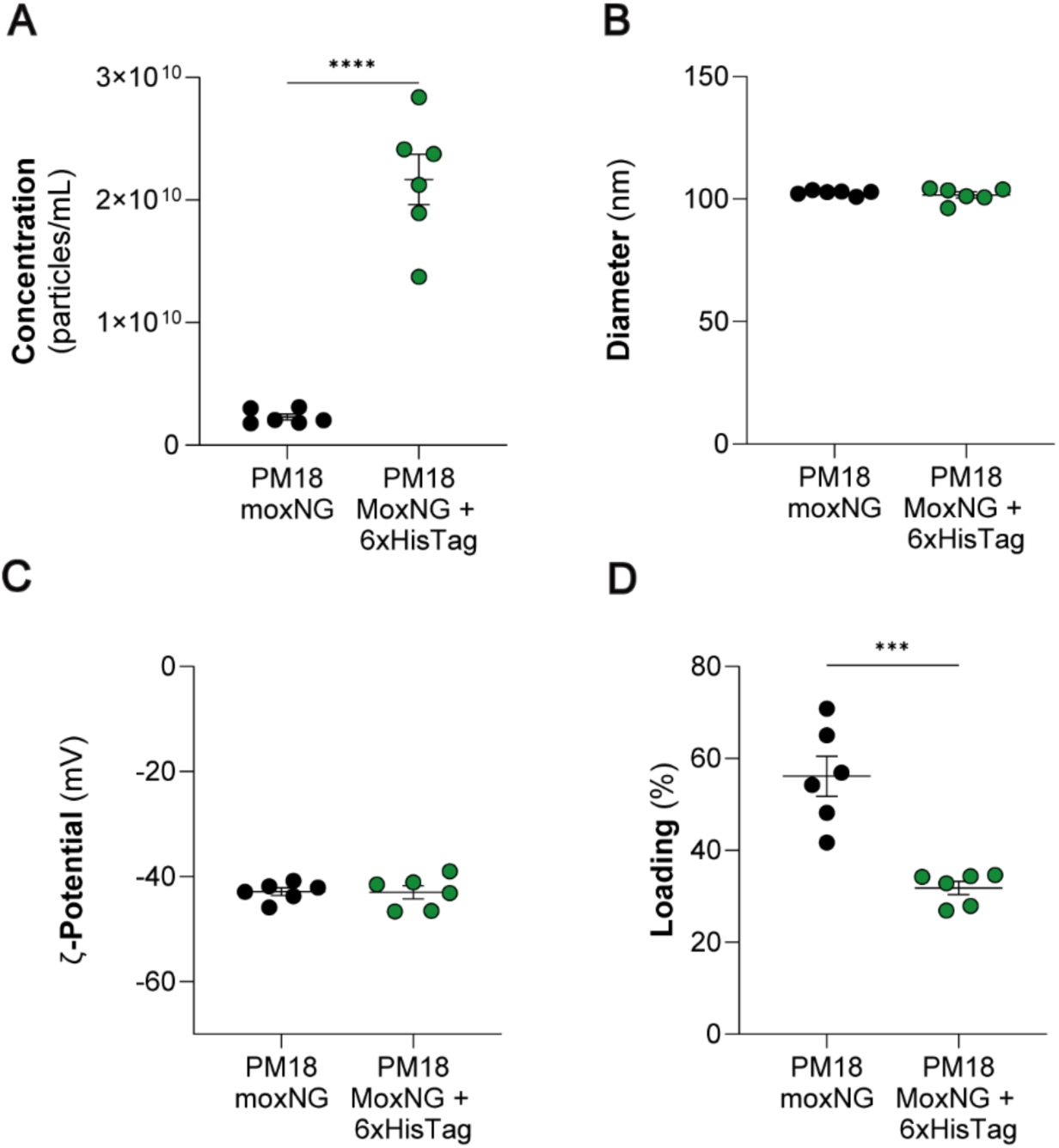
Comparison of PM18 moxNG bEVs and PM18 moxNG + 6xHisTag bEVs. The addition of a HisTag allows for quantification of moxNG via Western. (**A**) A higher concentration of bEVs was seen in PM18 moxNG + 6xHisTag bEVs (2.17 ± 0.21 × 10^10^ bEVs/mL) compared to PM18 moxNG (2.31 ± 0.24 × 10^9^ bEVs/mL, *p* < 0.0001). No difference was seen in (**B**) size or (**C**) ζ-potential. (**D**) PM18 moxNG bEVs had higher loading efficiency (56.15 ± 4.37%) compared to PM18 moxNG + 6xHisTag bEVs (31.81 ± 1.43%, *p* = 0.0003). Unpaired T-tests were used to determine significance. (\**p* ≤ 0.05, \*\**p* ≤ 0.01, \*\*\**p* ≤ 0.001, and \*\*\*\**p* ≤ 0.0001).

**Supplemental Figure 5:**
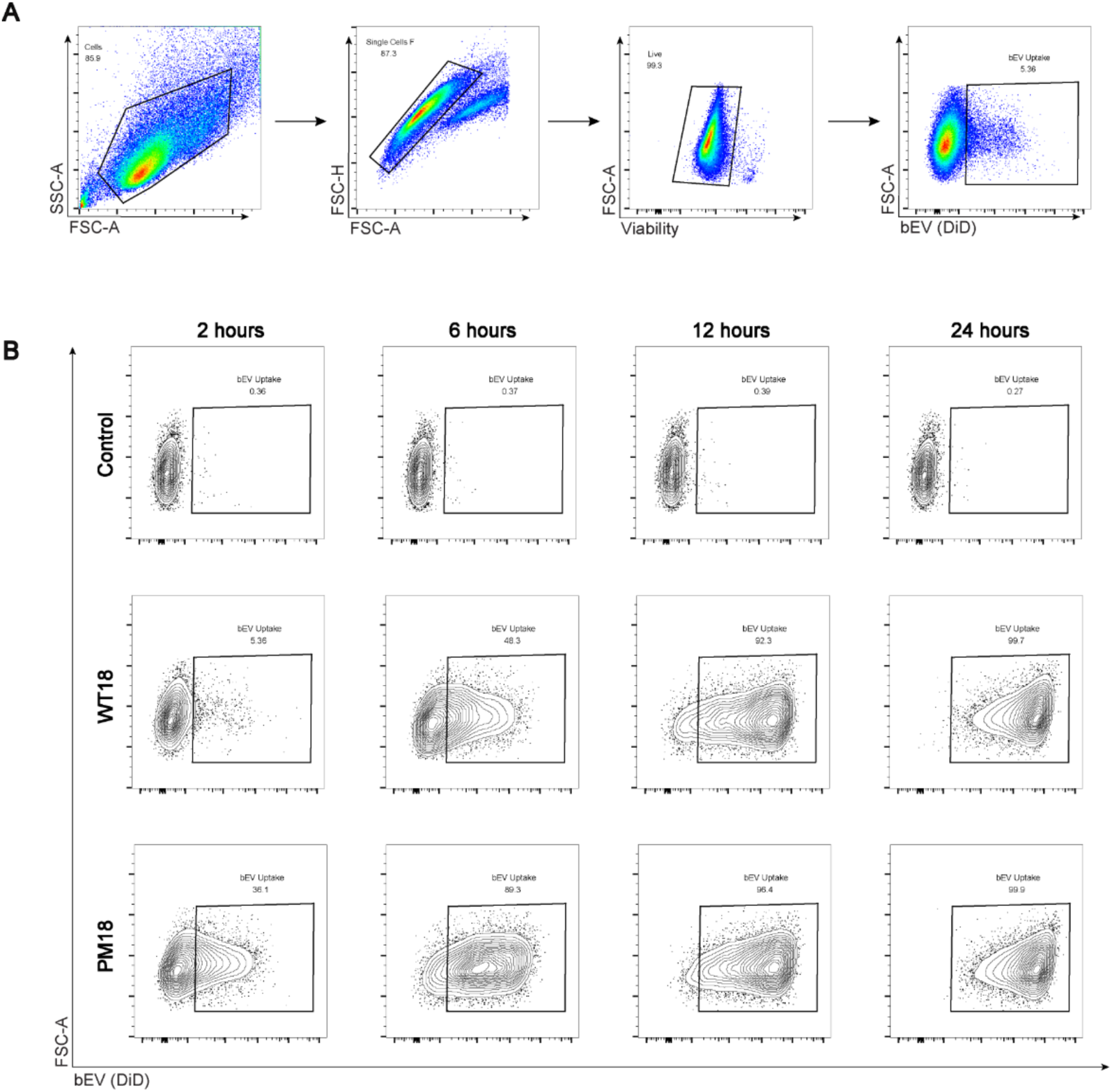
Determination of bEV uptake in vaginal epithelial cells via flow cytometry experiments. (**A**) Representative plots shown to illustrate gating. Flow cytometry gating removed cell debris, aggregates, and dead cells. Fluorescence intensity was used to determine gating for bEV+ (DiD+) cells. (**B**) Representative plots of conditions at each time point. Negative controls show limited residual background compared to treatment groups. Both treatment groups show vaginal epithelial cells increase in fluorescence with time, corresponding to an increase in uptake.

**Supplemental Figure 6:**
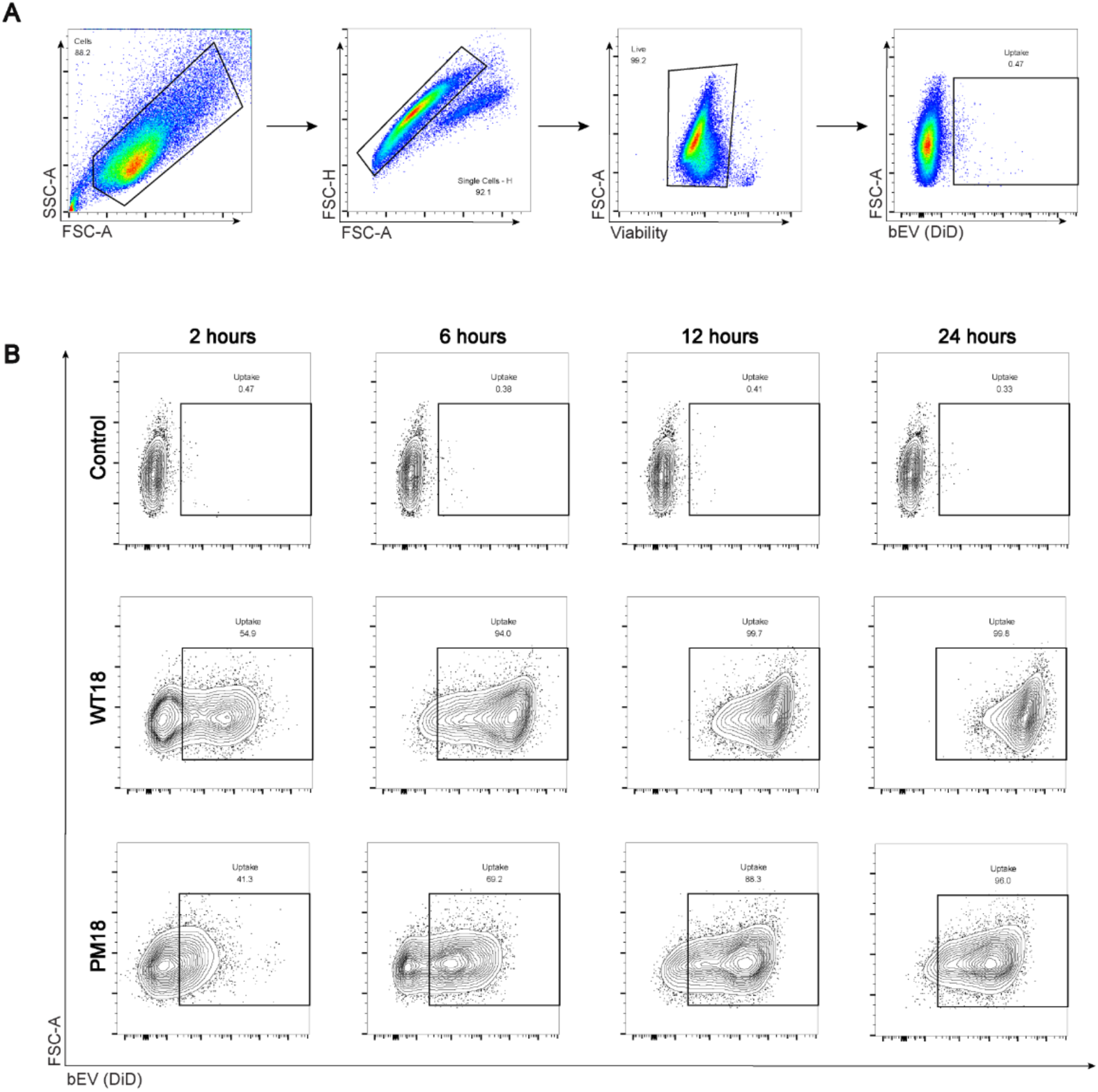
Determination of bEV uptake in endocervical cells via flow cytometry experiments. (**A**) Representative plots shown to illustrate gating. Flow cytometry gating removed cell debris, aggregates, and dead cells. Fluorescence intensity was used to determine gating for bEV+ (DiD+) cells. (**B**) Representative plots of conditions at each time point. Negative controls show limited residual background compared to treatment groups. Both treatment groups show endocervical cells increase in fluorescence with time, corresponding to an increase in uptake.

## References

1. Steinberg JR, Magnani CJ, Turner BE, Weeks BT, Young AMP, Lu CF, Zhang N, Richardson MT, Fitzgerald AC, Mekonnen Z, Redman T, Adetunji M, Martin SA, Anderson JN, Chan KS, Milad MP. Early Discontinuation, Results Reporting, and Publication of Gynecology Clinical Trials From 2007 to 2020. Obstetrics & Gynecology. 2022;139(5).

2. Mirin AA. Gender Disparity in the Funding of Diseases by the U.S. National Institutes of Health. Journal of Women’s Health. 2020;30(7):956–63. doi: 10.1089/jwh.2020.8682.

3. Mundekkad D, Cho WC. Nanoparticles in Clinical Translation for Cancer Therapy. International Journal of Molecular Sciences. 2022;23(3):1685. PubMed PMID: doi:10.3390/ijms23031685.

4. Gowd V, Ahmad A, Tarique M, Suhail M, Zughaibi TA, Tabrez S, Khan R. Advancement of cancer immunotherapy using nanoparticles-based nanomedicine. Seminars in Cancer Biology. 2022;86:624–44. doi: 10.1016/j.semcancer.2022.03.026.

5. Kirtane AR, Verma M, Karandikar P, Furin J, Langer R, Traverso G. Nanotechnology approaches for global infectious diseases. Nature Nanotechnology. 2021;16(4):369–84. doi: 10.1038/s41565-021-00866-8.

6. Pilkington EH, Suys EJA, Trevaskis NL, Wheatley AK, Zukancic D, Algarni A, Al-Wassiti H, Davis TP, Pouton CW, Kent SJ, Truong NP. From influenza to COVID-19: Lipid nanoparticle mRNA vaccines at the frontiers of infectious diseases. Acta Biomaterialia. 2021;131:16–40. doi: 10.1016/j.actbio.2021.06.023.

7. Butkovich N, Li E, Ramirez A, Burkhardt AM, Wang S-W. Advancements in protein nanoparticle vaccine platforms to combat infectious disease. WIREs Nanomedicine and Nanobiotechnology. 2021;13(3):e1681. doi: 10.1002/wnan.1681.

8. Lu Q, Kou D, Lou S, Ashrafizadeh M, Aref AR, Canadas I, Tian Y, Niu X, Wang Y, Torabian P, Wang L, Sethi G, Tergaonkar V, Tay F, Yuan Z, Han P. Nanoparticles in tumor microenvironment remodeling and cancer immunotherapy. Journal of Hematology & Oncology. 2024;17(1):16. doi: 10.1186/s13045-024-01535-8.

9. Shapiro RL, Bethiana T, Carter DM, Ortiz J, DeLong K, Anders N, Numan TA, Duggan E, Zierden HC, Ensign LM. Locally administered nanosuspension increases delivery of estradiol for the treatment of vaginal atrophy in mice. Drug Delivery and Translational Research. 2025;15(2):609–20. doi: 10.1007/s13346-024-01618-6.

10. Zhang T, Li D, Wang Y, Zhang C, Yang W, Gao G. Delivering umbilical cord mesenchymal stem cell exosomes through hydrogel ameliorates vaginal atrophy in ovariectomized rats. Aging (Albany NY). 2023;15(23):14292–305. Epub 20231206. doi: 10.18632/aging.205302. PubMed PMID: 38059876; PMCID: PMC10756086.

11. Zierden HC, Ortiz JI, DeLong K, Yu J, Li G, Dimitrion P, Bensouda S, Laney V, Bailey A, Anders NM, Scardina M, Mahendroo M, Mesiano S, Burd I, Wagner G, Hanes J, Ensign LM. Enhanced drug delivery to the reproductive tract using nanomedicine reveals therapeutic options for prevention of preterm birth. Science Translational Medicine. 2021;13(576):eabc6245. doi: 10.1126/scitranslmed.abc6245.

12. Yuan Y-G, Zhang S, Hwang J-Y, Kong I-K. Silver Nanoparticles Potentiates Cytotoxicity and Apoptotic Potential of Camptothecin in Human Cervical Cancer Cells. Oxidative Medicine and Cellular Longevity. 2018;2018(1):6121328. doi: 10.1155/2018/6121328.

13. Luo C-l, Liu Y-q, Wang P, Song C-h, Wang K-j, Dai L-p, Zhang J-y, Ye H. The effect of quercetin nanoparticle on cervical cancer progression by inducing apoptosis, autophagy and anti-proliferation via JAK2 suppression. Biomedicine & Pharmacotherapy. 2016;82:595–605. doi: 10.1016/j.biopha.2016.05.029.

14. Khan MA, Zafaryab M, Mehdi SH, Ahmad I, Rizvi MMA. Characterization and anti-proliferative activity of curcumin loaded chitosan nanoparticles in cervical cancer. International Journal of Biological Macromolecules. 2016;93:242–53. doi: 10.1016/j.ijbiomac.2016.08.050.

15. Ensign LM, Tang BC, Wang Y-Y, Tse TA, Hoen T, Cone R, Hanes J. Mucus-Penetrating Nanoparticles for Vaginal Drug Delivery Protect Against Herpes Simplex Virus. Science Translational Medicine. 2012;4(138):138ra79-ra79. doi: doi:10.1126/scitranslmed.3003453.

16. Mohammed Fayaz A, Zhujun A, Morkattu G, Liyu C, Xianzhong X, T. KP, and Yao X. Inactivation of microbial infectiousness by silver nanoparticles-coated condom: a new approach to inhibit HIV- and HSV-transmitted infection. International Journal of Nanomedicine. 2012;7(null):5007–18. doi: 10.2147/IJN.S34973.

17. Kirian RD, Steinman D, Jewell CM, Zierden HC. Extracellular vesicles as carriers of mRNA: Opportunities and challenges in diagnosis and treatment. Theranostics. 2024;14(5):2265–89. Epub 20240311. doi: 10.7150/thno.93115. PubMed PMID: 38505610; PMCID: PMC10945352.

18. Liu H, Geng Z, Su J. Engineered mammalian and bacterial extracellular vesicles as promising nanocarriers for targeted therapy. Extracell Vesicles Circ Nucl Acids. 2022;3(2):63–86. Epub 20220413. doi: 10.20517/evcna.2022.04. PubMed PMID: 39698442; PMCID: PMC11648430.

19. Liu H, Li M, Zhang T, Liu X, Zhang H, Geng Z, Su J. Engineered bacterial extracellular vesicles for osteoporosis therapy. Chemical Engineering Journal. 2022;450:138309. doi: 10.1016/j.cej.2022.138309.

20. Lin Y, Jingyu W, Fan B, Ruyi Z, Junhui W, Yubing W, Mei H, Yiyi H, Lei Z, Qian W, and Hu X. Bacterial extracellular vesicles in the initiation, progression and treatment of atherosclerosis. Gut Microbes. 2025;17(1):2452229. doi: 10.1080/19490976.2025.2452229.

21. Choi Y, Park HS, Kim YK. Bacterial Extracellular Vesicles: A Candidate Molecule for the Diagnosis and Treatment of Allergic Diseases. Allergy Asthma Immunol Res. 2023;15(3):279–89. doi: 10.4168/aair.2023.15.3.279. PubMed PMID: 37188485; PMCID: PMC10186123.

22. Ji N, Wang F, Wang M, Zhang W, Liu H, Su J. Engineered bacterial extracellular vesicles for central nervous system diseases. Journal of Controlled Release. 2023;364:46–60. doi: 10.1016/j.jconrel.2023.10.027.

23. Moore KA, Petersen AP, Zierden HC. Microorganism-derived extracellular vesicles: emerging contributors to female reproductive health. Nanoscale. 2024;16(17):8216–35. Epub 20240502. doi: 10.1039/d3nr05524h. PubMed PMID: 38572613.

24. Ñahui Palomino RA, Vanpouille C, Costantini PE, Margolis L. Microbiota–host communications: Bacterial extracellular vesicles as a common language. PLOS Pathogens. 2021;17(5):e1009508. doi: 10.1371/journal.ppat.1009508.

25. Nie X, Li Q, Chen X, Onyango S, Xie J, Nie S. Bacterial extracellular vesicles: Vital contributors to physiology from bacteria to host. Microbiological Research. 2024;284:127733. doi: 10.1016/j.micres.2024.127733.

26. Lee W-H, Choi H-I, Hong S-W, Kim K-s, Gho YS, Jeon SG. Vaccination with Klebsiella pneumoniae-derived extracellular vesicles protects against bacteria-induced lethality via both humoral and cellular immunity. Experimental & Molecular Medicine. 2015;47(9):e183-e. doi: 10.1038/emm.2015.59.

27. Jiang L, Driedonks TAP, Jong WSP, Dhakal S, Bart van den Berg van Saparoea H, Sitaras I, Zhou R, Caputo C, Littlefield K, Lowman M, Chen M, Lima G, Gololobova O, Smith B, Mahairaki V, Riley Richardson M, Mulka KR, Lane AP, Klein SL, Pekosz A, Brayton C, Mankowski JL, Luirink J, Villano JS, Witwer KW. A bacterial extracellular vesicle-based intranasal vaccine against SARS-CoV-2 protects against disease and elicits neutralizing antibodies to wild-type and Delta variants. Journal of Extracellular Vesicles. 2022;11(3):e12192. doi: 10.1002/jev2.12192.

28. Schoch CL, Ciufo S, Domrachev M, Hotton CL, Kannan S, Khovanskaya R, Leipe D, McVeigh R, O’Neill K, Robbertse B, Sharma S, Soussov V, Sullivan JP, Sun L, Turner S, Karsch-Mizrachi I. NCBI Taxonomy: a comprehensive update on curation, resources and tools. Database. 2020;2020:baaa062. doi: 10.1093/database/baaa062.

29. Molinari S, Tesoriero RF, Li D, Sridhar S, Cai R, Soman J, Ryan KR, Ashby PD, Ajo-Franklin CM. A de novo matrix for macroscopic living materials from bacteria. Nature Communications. 2022;13(1):5544. doi: 10.1038/s41467-022-33191-2.

30. Sarra A, Celluzzi A, Bruno SP, Ricci C, Sennato S, Ortore MG, Casciardi S, Del Chierico F, Postorino P, Bordi F, Masotti A. Biophysical Characterization of Membrane Phase Transition Profiles for the Discrimination of Outer Membrane Vesicles (OMVs) From Escherichia coli Grown at Different Temperatures. Frontiers in Microbiology. 2020;Volume 11 - 2020. doi: 10.3389/fmicb.2020.00290.

31. Steinman D, Kirian RD, Zierden HC. Multiple Particle Tracking: A Method for Probing Biologically Relevant Mobility of Bacterial Extracellular Vesicles. In: Jay S, Pirolli N, editors. Bacterial Extracellular Vesicles: Methods and Protocols. New York, NY: Springer US; 2024. p. 137–52.

32. Pradhan S, Weiss Alison A. Probiotic Properties of Escherichia coli Nissle in Human Intestinal Organoids. mBio. 2020;11(4):10.1128/mbio.01470-20. doi: 10.1128/mbio.01470-20.

33. Schwechheimer C, Sullivan CJ, Kuehn MJ. Envelope Control of Outer Membrane Vesicle Production in Gram-Negative Bacteria. Biochemistry. 2013;52(18):3031–40. doi: 10.1021/bi400164t.

34. Bonnington KE, Kuehn MJ. Protein selection and export via outer membrane vesicles. Biochimica et Biophysica Acta (BBA) - Molecular Cell Research. 2014;1843(8):1612–9. doi: 10.1016/j.bbamcr.2013.12.011.

35. Beveridge TJ. Structures of gram-negative cell walls and their derived membrane vesicles. J Bacteriol. 1999;181(16):4725–33. doi: 10.1128/jb.181.16.4725-4733.1999. PubMed PMID: 10438737; PMCID: PMC93954.

36. Wo J, Lv Z-Y, Sun J-N, Tang H, Qi N, Ye B-C. Engineering probiotic-derived outer membrane vesicles as functional vaccine carriers to enhance immunity against SARS-CoV-2. iScience. 2023;26(1):105772. doi: 10.1016/j.isci.2022.105772.

37. Fantappiè L, Micaela dS, Emiliano C, Filippo C, Giuliano B, Olivier J, Immaculada M, and Grandi G. Antibody-mediated immunity induced by engineered Escherichia coli OMVs carrying heterologous antigens in their lumen. Journal of Extracellular Vesicles. 2014;3(1):24015. doi: 10.3402/jev.v3.24015.

38. Osgerby A, Overton TW. Approaches for high-throughput quantification of periplasmic recombinant proteins. New Biotechnology. 2023;77:149–60. doi: 10.1016/j.nbt.2023.09.003.

39. Karyolaimos A, Dolata KM, Antelo-Varela M, Mestre Borras A, Elfageih R, Sievers S, Becher D, Riedel K, de Gier J-W. Escherichia coli Can Adapt Its Protein Translocation Machinery for Enhanced Periplasmic Recombinant Protein Production. Frontiers in Bioengineering and Biotechnology. 2020;Volume 7 - 2019. doi: 10.3389/fbioe.2019.00465.

40. Bhatwa A, Wang W, Hassan YI, Abraham N, Li X-Z, Zhou T. Challenges Associated With the Formation of Recombinant Protein Inclusion Bodies in Escherichia coli and Strategies to Address Them for Industrial Applications. Frontiers in Bioengineering and Biotechnology. 2021;Volume 9 - 2021. doi: 10.3389/fbioe.2021.630551.

41. Théry C, Witwer KW, Aikawa E, Alcaraz MJ, Anderson JD, Andriantsitohaina R, Antoniou A, Arab T, Archer F, Atkin-Smith GK, Ayre DC, Bach J-M, Bachurski D, Baharvand H, Balaj L, Baldacchino S, Bauer NN, Baxter AA, Bebawy M, Beckham C, Bedina Zavec A, Benmoussa A, Berardi AC, Bergese P, Bielska E, Blenkiron C, Bobis-Wozowicz S, Boilard E, Boireau W, Bongiovanni A, Borràs FE, Bosch S, Boulanger CM, Breakefield X, Breglio AM, Brennan MÁ, Brigstock DR, Brisson A, Broekman ML, Bromberg JF, Bryl-Górecka P, Buch S, Buck AH, Burger D, Busatto S, Buschmann D, Bussolati B, Buzás EI, Byrd JB, Camussi G, Carter DR, Caruso S, Chamley LW, Chang Y-T, Chen C, Chen S, Cheng L, Chin AR, Clayton A, Clerici SP, Cocks A, Cocucci E, Coffey RJ, Cordeiro-da-Silva A, Couch Y, Coumans FA, Coyle B, Crescitelli R, Criado MF, D’Souza-Schorey C, Das S, Datta Chaudhuri A, de Candia P, De Santana Junior EF, De Wever O, del Portillo HA, Demaret T, Deville S, Devitt A, Dhondt B, Di Vizio D, Dieterich LC, Dolo V, Dominguez Rubio AP, Dominici M, Dourado MR, Driedonks TA, Duarte FV, Duncan HM, Eichenberger RM, Ekström K, EL Andaloussi S, Elie-Caille C, Erdbrügger U, Falcón-Pérez JM, Fatima F, Fish JE, Flores-Bellver M, Försönits A, Frelet-Barrand A, Fricke F, Fuhrmann G, Gabrielsson S, Gámez-Valero A, Gardiner C, Gärtner K, Gaudin R, Gho YS, Giebel B, Gilbert C, Gimona M, Giusti I, Goberdhan DC, Görgens A, Gorski SM, Greening DW, Gross JC, Gualerzi A, Gupta GN, Gustafson D, Handberg A, Haraszti RA, Harrison P, Hegyesi H, Hendrix A, Hill AF, Hochberg FH, Hoffmann KF, Holder B, Holthofer H, Hosseinkhani B, Hu G, Huang Y, Huber V, Hunt S, Ibrahim AG-E, Ikezu T, Inal JM, Isin M, Ivanova A, Jackson HK, Jacobsen S, Jay SM, Jayachandran M, Jenster G, Jiang L, Johnson SM, Jones JC, Jong A, Jovanovic-Talisman T, Jung S, Kalluri R, Kano S-i, Kaur S, Kawamura Y, Keller ET, Khamari D, Khomyakova E, Khvorova A, Kierulf P, Kim KP, Kislinger T, Klingeborn M, Klinke II DJ, Kornek M, Kosanović MM, Kovács ÁF, Krämer-Albers E-M, Krasemann S, Krause M, Kurochkin IV, Kusuma GD, Kuypers S, Laitinen S, Langevin SM, Languino LR, Lannigan J, Lässer C, Laurent LC, Lavieu G, Lázaro-Ibáñez E, Le Lay S, Lee M-S, Lee YXF, Lemos DS, Lenassi M, Leszczynska A, Li IT, Liao K, Libregts SF, Ligeti E, Lim R, Lim SK, Linē A, Linnemannstöns K, Llorente A, Lombard CA, Lorenowicz MJ, Lörincz ÁM, Lötvall J, Lovett J, Lowry MC, Loyer X, Lu Q, Lukomska B, Lunavat TR, Maas SL, Malhi H, Marcilla A, Mariani J, Mariscal J, Martens-Uzunova ES, Martin-Jaular L, Martinez MC, Martins VR, Mathieu M, Mathivanan S, Maugeri M, McGinnis LK, McVey MJ, Meckes Jr DG, Meehan KL, Mertens I, Minciacchi VR, Möller A, Møller Jørgensen M, Morales-Kastresana A, Morhayim J, Mullier F, Muraca M, Musante L, Mussack V, Muth DC, Myburgh KH, Najrana T, Nawaz M, Nazarenko I, Nejsum P, Neri C, Neri T, Nieuwland R, Nimrichter L, Nolan JP, Nolte-’t Hoen EN, Noren Hooten N, O’Driscoll L, O’Grady T, O’Loghlen A, Ochiya T, Olivier M, Ortiz A, Ortiz LA, Osteikoetxea X, Østergaard O, Ostrowski M, Park J, Pegtel DM, Peinado H, Perut F, Pfaffl MW, Phinney DG, Pieters BC, Pink RC, Pisetsky DS, Pogge von Strandmann E, Polakovicova I, Poon IK, Powell BH, Prada I, Pulliam L, Quesenberry P, Radeghieri A, Raffai RL, Raimondo S, Rak J, Ramirez MI, Raposo G, Rayyan MS, Regev-Rudzki N, Ricklefs FL, Robbins PD, Roberts DD, Rodrigues SC, Rohde E, Rome S, Rouschop KM, Rughetti A, Russell AE, Saá P, Sahoo S, Salas-Huenuleo E, Sánchez C, Saugstad JA, Saul MJ, Schiffelers RM, Schneider R, Schøyen TH, Scott A, Shahaj E, Sharma S, Shatnyeva O, Shekari F, Shelke GV, Shetty AK, Shiba K, Siljander PR-M, Silva AM, Skowronek A, Snyder II OL, Soares RP, Sódar BW, Soekmadji C, Sotillo J, Stahl PD, Stoorvogel W, Stott SL, Strasser EF, Swift S, Tahara H, Tewari M, Timms K, Tiwari S, Tixeira R, Tkach M, Toh WS, Tomasini R, Torrecilhas AC, Tosar JP, Toxavidis V, Urbanelli L, Vader P, van Balkom BW, van der Grein SG, Van Deun J, van Herwijnen MJ, Van Keuren-Jensen K, van Niel G, van Royen ME, van Wijnen AJ, Vasconcelos MH, Vechetti Jr IJ, Veit TD, Vella LJ, Velot É, Verweij FJ, Vestad B, Viñas JL, Visnovitz T, Vukman KV, Wahlgren J, Watson DC, Wauben MH, Weaver A, Webber JP, Weber V, Wehman AM, Weiss DJ, Welsh JA, Wendt S, Wheelock AM, Wiener Z, Witte L, Wolfram J, Xagorari A, Xander P, Xu J, Yan X, Yáñez-Mó M, Yin H, Yuana Y, Zappulli V, Zarubova J, Žėkas V, Zhang J-y, Zhao Z, Zheng L, Zheutlin AR, Zickler AM, Zimmermann P, Zivkovic AM, Zocco D, Zuba-Surma EK. Minimal information for studies of extracellular vesicles 2018 (MISEV2018): A position statement of the International Society for Extracellular Vesicles and update of the MISEV2014 guidelines. J Extracell Vesicles. 2018;7(1):1535750. doi: 10.1080/20013078.2018.1535750.

42. Infertility and Impaired Fecundity in Women and Men in the United States, 2015–2019, (2024).

43. Riestenberg C, Jagasia A, Markovic D, Buyalos RP, Azziz R. Health Care-Related Economic Burden of Polycystic Ovary Syndrome in the United States: Pregnancy-Related and Long-Term Health Consequences. The Journal of Clinical Endocrinology & Metabolism. 2021;107(2):575–85. doi: 10.1210/clinem/dgab613.

44. Huang S-H, Hsu H-C, Lee T-F, Fan H-M, Tseng C-W, Chen IH, Shen H, Lee C-Y, Tai H-T, Hsu H-M, Hung C-C. Prevalence, Associated Factors, and Appropriateness of Empirical Treatment of Trichomoniasis, Bacterial Vaginosis, and Vulvovaginal Candidiasis among Women with Vaginitis. Microbiology Spectrum. 2023;11(3):e00161–23. doi: 10.1128/spectrum.00161-23.

45. Simoens S, Dunselman G, Dirksen C, Hummelshoj L, Bokor A, Brandes I, Brodszky V, Canis M, Colombo GL, DeLeire T, Falcone T, Graham B, Halis G, Horne A, Kanj O, Kjer JJ, Kristensen J, Lebovic D, Mueller M, Vigano P, Wullschleger M, D’Hooghe T. The burden of endometriosis: costs and quality of life of women with endometriosis and treated in referral centres. Human Reproduction. 2012;27(5):1292–9. doi: 10.1093/humrep/des073.

46. Mercuri ND, Cox BJ. The need for more research into reproductive health and disease. Elife. 2022;11. Epub 20221213. doi: 10.7554/eLife.75061. PubMed PMID: 36511240; PMCID: PMC9771341.

47. Guo J, Huang Z, Wang Q, Wang M, Ming Y, Chen W, Huang Y, Tang Z, Huang M, Liu H, Jia B. Opportunities and challenges of bacterial extracellular vesicles in regenerative medicine. Journal of Nanobiotechnology. 2025;23(1):4. doi: 10.1186/s12951-024-02935-1.

48. Liu C, Yazdani N, Moran CS, Salomon C, Seneviratne CJ, Ivanovski S, Han P. Unveiling clinical applications of bacterial extracellular vesicles as natural nanomaterials in disease diagnosis and therapeutics. Acta Biomaterialia. 2024;180:18–45. doi: 10.1016/j.actbio.2024.04.022.

49. Hosseini-Giv N, Basas A, Hicks C, El-Omar E, El-Assaad F, Hosseini-Beheshti E. Bacterial extracellular vesicles and their novel therapeutic applications in health and cancer. Frontiers in Cellular and Infection Microbiology. 2022;Volume 12 - 2022. doi: 10.3389/fcimb.2022.962216.

50. Srikrishna S, Cardozo L. The vagina as a route for drug delivery: a review. International Urogynecology Journal. 2013;24(4):537–43. doi: 10.1007/s00192-012-2009-3.

51. Subi MTM, Selvasudha N, Vasanthi HR. Vaginal drug delivery system: A promising route of drug administration for local and systemic diseases. Drug Discovery Today. 2024;29(6):104012. doi: 10.1016/j.drudis.2024.104012.

52. De Ziegler D, Bulletti C, De Monstier B, Jaaskelainen AS. The first uterine pass effect. Ann N Y Acad Sci. 1997;828:291–9. Epub 1997/11/05. PubMed PMID: 9329850.

53. Carson L, Merkatz R, Martinelli E, Boyd P, Variano B, Sallent T, Malcolm RK. The Vaginal Microbiota, Bacterial Biofilms and Polymeric Drug-Releasing Vaginal Rings. Pharmaceutics. 2021;13(5):751. PubMed PMID: doi:10.3390/pharmaceutics13050751.

54. Zierden HC, DeLong K, Zulfiqar F, Ortiz JO, Laney V, Bensouda S, Hernández N, Hoang TM, Lai SK, Hanes J, Burke AE, Ensign LM. Cervicovaginal mucus barrier properties during pregnancy are impacted by the vaginal microbiome. Frontiers in Cellular and Infection Microbiology. 2023;13. doi: 10.3389/fcimb.2023.1015625.

55. Zierden HC, Josyula A, Shapiro RL, Hsueh HT, Hanes J, Ensign LM. Avoiding a Sticky Situation: Bypassing the Mucus Barrier for Improved Local Drug Delivery. Trends in Molecular Medicine. 2021;27(5):436–50. doi: 10.1016/j.molmed.2020.12.001.

56. Dedeloudi A, Siamidi A, Pavlou P, Vlachou M. Recent Advances in the Excipients Used in Modified Release Vaginal Formulations. Materials. 2022;15(1):327. PubMed PMID: doi:10.3390/ma15010327.

57. Joseph A, Anton L, Guan Y, Ferguson B, Mirro I, Meng N, France M, Ravel J, Elovitz MA. Extracellular vesicles from vaginal Gardnerella vaginalis and Mobiluncus mulieris contain distinct proteomic cargo and induce inflammatory pathways. npj Biofilms and Microbiomes. 2024;10(1):28. doi: 10.1038/s41522-024-00502-y.

58. Shishpal P, Kasarpalkar N, Singh D, Bhor VM. Characterization of Gardnerella vaginalis membrane vesicles reveals a role in inducing cytotoxicity in vaginal epithelial cells. Anaerobe. 2020;61:102090. Epub 2019/08/24. doi: 10.1016/j.anaerobe.2019.102090. PubMed PMID: 31442559.

59. Shi J, Ma D, Gao S, Long F, Wang X, Pu X, Cannon RD, Han T-L. Probiotic Escherichia coli Nissle 1917-derived outer membrane vesicles modulate the intestinal microbiome and host gut-liver metabolome in obese and diabetic mice. Frontiers in Microbiology. 2023;Volume 14 - 2023. doi: 10.3389/fmicb.2023.1219763.

60. Li N, Xin H, Deng K. Separation and Characterization of Heterogeneity Among Various Sizes of Outer Membrane Vesicles Derived from the Probiotic Escherichia coli Nissle 1917. Membranes. 2025;15(5):141. PubMed PMID: doi:10.3390/membranes15050141.

61. Sawabe T, Ojima Y, Nakagawa M, Sawada T, Tahara YO, Miyata M, Azuma M. Construction and characterization of a hypervesiculation strain of Escherichia coli Nissle 1917. PLOS ONE. 2024;19(4):e0301613. doi: 10.1371/journal.pone.0301613.

62. Wang X, Liu S, Guan Y, Ding J, Ma C, Xie Z. Vaginal drug delivery approaches for localized management of cervical cancer. Advanced Drug Delivery Reviews. 2021;174:114–26. doi: 10.1016/j.addr.2021.04.009.

63. El-Hammadi MM, Arias JL. Chapter 11 - Nanomedicine for vaginal drug delivery. In: Kesharwani P, Taurin S, Greish K, editors. Theory and Applications of Nonparenteral Nanomedicines: Academic Press; 2021. p. 235-57.

