## Supplementary Information for "Bacterial extracellular vesicles as a tunable platform for vaginal drug delivery"

### Supplementary Material

#### Table 1: Sequences for relevant proteins

| Protein Complex | Sequence |
|---|---|
| moxNeonGreen | ATGgtttcgaaaggtgaggaggataacatggcgtcgctgcctgctacacatgaactgcatattttcggcagcatcaacggagttgatttcgaca<br>tggtgggccaaggcaccggcaacccgaacgacggctatgaagaattgaatttgaaatctaccaagggcgacttgcaatttagtccgtggattct<br>ggtaccgcatattggttatggttttcatcagtatctgccctacccagacggaatgagcccatttcaggcggctatggtggatggcagtggctatcag<br>gtccatcgcaccatgcaattcgaggatggggcgagcctgaccgtgaattatcggtacacttacgaaggttcacatattaaaggtgaagcacaa<br>gttaaaggcaccggcttcccggcggatggcccagtgatgaccaacagcctgacggcggcggactggtcccgtagcaaaaagacgtatccc<br>aatgataaaacaattatttctacctttaaatggagttacaccacgggtaacggtaaacgctaccgtagcacggctcgcaccacatacacctttgc<br>gaaacccatggccgcgaactatctgaaaaatcagccgatgtatgtgtttcgcaaaacggaactgaaacattcgaaaactgaactcaatttcaa<br>agagtggcaaaaagccttcaccgacgtaatgggcatggatgaactgtacaaa |
| OmpA-moxNeonGreen | ATGAAAAAGACAGCTATCGCGATTGCAGTGGCACTGGCTGGTTTCGCTACCGTAGCGCAGGCCG<br>CTCCGAAAGATatggtttcgaaaggtgaggaggataacatggcgtcgctgcctgctacacatgaactgcatattttcggcagcatcaacg<br>gagttgatttcgacatggtgggccaaggcaccggcaacccgaacgacggctatgaagaattgaatttgaaatctaccaagggcgacttgcaa<br>tttagtccgtggattctggtaccgcatattggttatggttttcatcagtatctgccctacccagacggaatgagcccatttcaggcggctatggtggat<br>ggcagtggctatcaggtccatcgcaccatgcaattcgaggatggggcgagcctgaccgtgaattatcggtacacttacgaaggttcacatatta<br>aaggtgaagcacaagttaaaggcaccggcttcccggcggatggcccagtgatgaccaacagcctgacggcggcggactggtcccgtagca<br>aaaagacgtatcccaatgataaaacaattatttctacctttaaatggagttacaccacgggtaacggtaaacgctaccgtagcacggctcgcac<br>cacatacacctttgcgaaacccatggccgcgaactatctgaaaaatcagccgatgtatgtgtttcgcaaaacggaactgaaacattcgaaaac<br>tgaactcaatttcaaagagtggcaaaaagccttcaccgacgtaatgggcatggatgaactgtacaaatga |

#### Table 2: Primers for relevant plasmids

| Plasmid | Forward sequence 5'-3' | Reverse sequence 3'-5' |
|---|---|---|
| pVHSM035 | CAGGTCTCGACCATCACCATCACtgaaagcttaattagctg | CTGGTCTCATGGTGATGCGATCCtttgtacagttcatccat |
| pRASM009 | TCGGTCTCGctATGgtttcgaaaggtgag | CTGGTCTCGATagttaatttctcctctttaatg |
| pRASM010 | CTGGTCTCGctATGgtttcgaaaggtgag | CTGGTCTCGATagttaatttctcctctttaatg |
| pVHSM045 | GAGGTCTCCATCACCATCACtaactcgaggaattcgaa | GAGGTCTCCTGCCCTGAAAATACAGGTT |

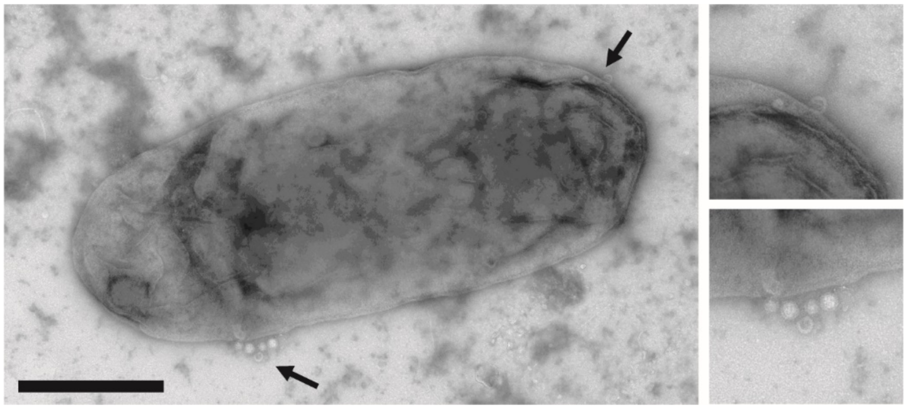

**Supplemental Figure 1: Verification of bEV production by *Escherichia coli* Nissle 1917.** Diluted cultures were negatively stained via uranyl acetate and imaged via transmission electron microscopy at the University of Maryland, Laboratory for Biological Ultrastructure. Arrows denote regions of nanoparticles determined to be bEVs. Scale bar denotes 500 nm.

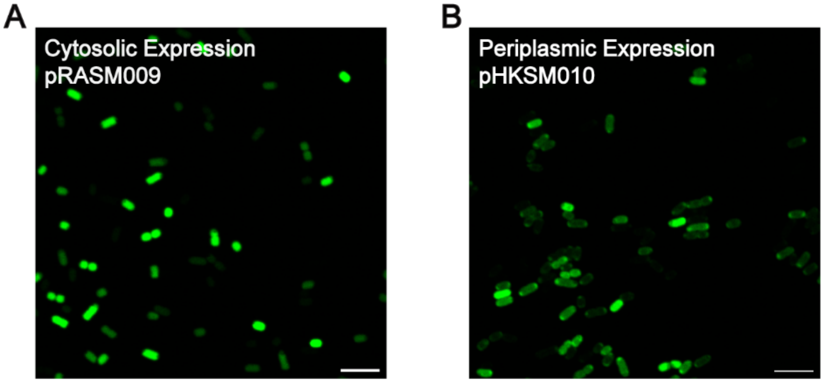


**Supplemental Figure 2: Confocal microscopy confirms moxNG production.** Production of moxNG in EcN in the (**A**) cytosol and (**A**) periplasmic was confirmed via confocal microscopy. Periplasmic localization is indicated by fluorescence restricted to the cell periphery. Scale bars denote 5 μm.

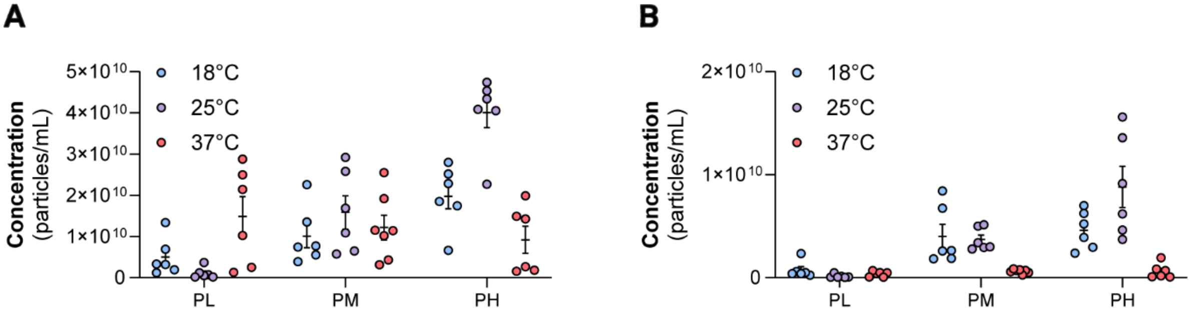


**Supplemental Figure 3: Original bEV concentration for loading optimization experiments.** Loading optimization of WT bacteria with periplasmic moxNG expression was determined at 3 different IPTG induction levels, low (1.56 nM; PL), medium (6.25 nM; PM) or high (25 nM; PH), and 3 different temperatures (18˚C, 25˚C, or 37˚C). The concentration of the (**A**) total population and (**B**) fluorescent population were determined via nanoparticle tracking analysis.

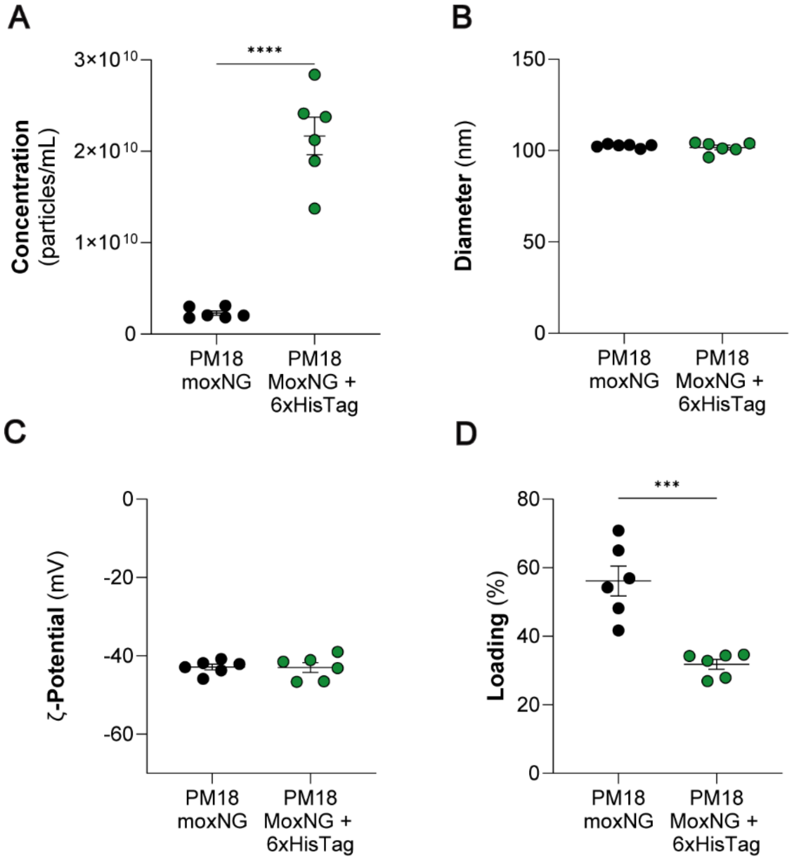


**Supplemental Figure 4: Comparison of PM18 moxNG bEVs and PM18 moxNG + 6xHisTag bEVs.** The addition of a HisTag allows for quantification of moxNG via Western. (**A**) A higher concentration of bEVs was seen in PM18 moxNG + 6xHisTag bEVs (2.17 ± 0.21 x $10^{10}$ bEVs/mL) compared to PM18 moxNG (2.31 ± 0.24 x $10^{9}$ bEVs/mL, $p < 0.0001$). No difference was seen in (**B**) size or (**C**) ζ-potential. (**D**) PM18 moxNG bEVs had higher loading efficiency (56.15 ± 4.37%) compared to PM18 moxNG + 6xHisTag bEVs (31.81 ± 1.43%, $p = 0.0003$). Unpaired T-tests were used to determine significance. (*$p \leq 0.05$, **$p \leq 0.01$, ***$p \leq 0.001$, and ****$p \leq 0.0001$).

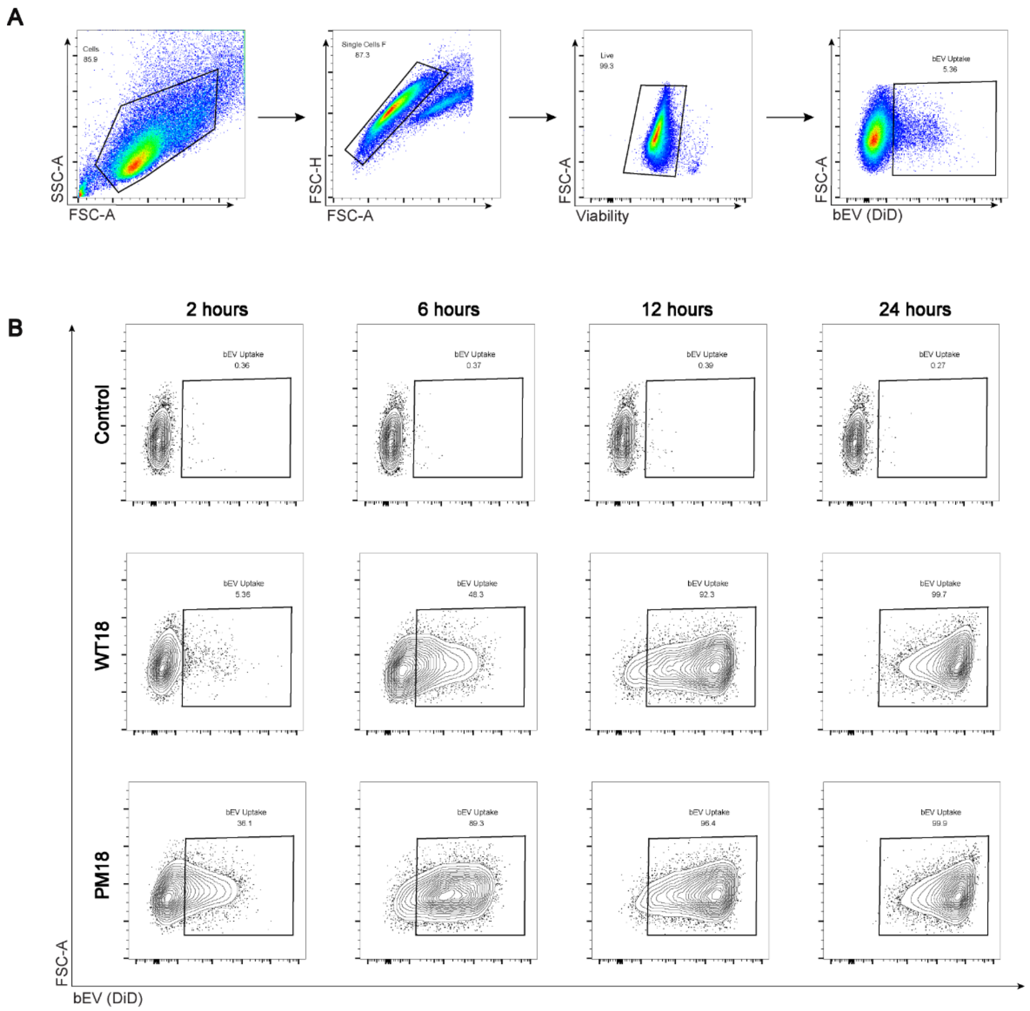


**Supplemental Figure 5: Determination of bEV uptake in vaginal epithelial cells via flow cytometry experiments.** (**A**) Representative plots shown to illustrate gating. Flow cytometry gating removed cell debris, aggregates, and dead cells. Fluorescence intensity was used to determine gating for bEV+ (DiD+) cells. (**B**) Representative plots of conditions at each time point. Negative controls show limited residual background compared to treatment groups. Both treatment groups show vaginal epithelial cells increase in fluorescence with time, corresponding to an increase in uptake.

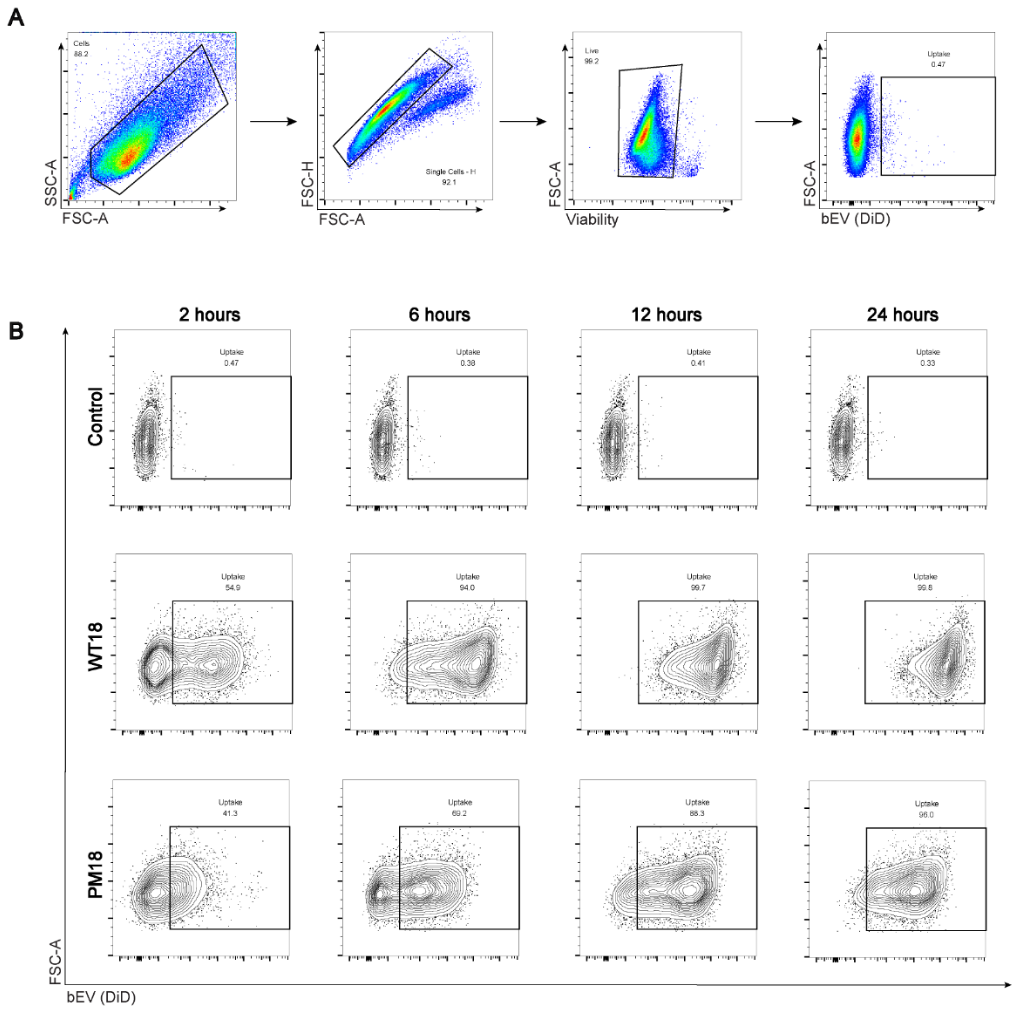


**Supplemental Figure 6: Determination of bEV uptake in endocervical cells via flow cytometry experiments.** (**A**) Representative plots shown to illustrate gating. Flow cytometry gating removed cell debris, aggregates, and dead cells. Fluorescence intensity was used to determine gating for bEV+ (DiD+) cells. (**B**) Representative plots of conditions at each time point. Negative controls show limited residual background compared to treatment groups. Both treatment groups show endocervical cells increase in fluorescence with time, corresponding to an increase in uptake.
